# Cervical Repetitive Magnetic Stimulation Enhances Respiratory Recovery by Modulating Neuronal Plasticity After Cervical Spinal Cord Injury

**DOI:** 10.64898/2026.03.31.715726

**Authors:** Wei Chen, Stéphane Vinit, Isabelle Vivodtzev

## Abstract

Cervical spinal cord injury (SCI) frequently leads to life-threatening respiratory insufficiency by disrupting descending phrenic pathways. There is growing interest in non-invasive neuromodulatory approaches to enhance plasticity of spared respiratory circuits. We investigated whether cervical repetitive magnetic stimulation (rMS) applied to the injured cervical spinal cord promotes ventilatory recovery in a preclinical mouse model. Adult mice received a unilateral C3 hemicontusion followed by either rMS or sham stimulation. We found that rMS-treated mice significantly improved recovery of tidal volume and minute ventilation at 21 days post injury(dpi) compared to sham controls under various breathing conditions (isoflurane anesthesia, poikilocapnic phase and hypercapnic challenge). Correspondingly, diaphragm EMG enhanced ipsilateral hemidiaphragm activity in ventral and medial regions, and even contralateral hemidiaphragm activity in its ventral part. This was associated with a marked attenuation of the inflammatory response at the cervical spinal cord level. Indeed, rMS lowered astroglial, fibrotic scarring, pro-inflammatory CD68-, Iba1- microglial/macrophage markers. Moreover, perineuronal net expression (WFA positive staining) is globally reduced in the ventral spinal horn, whereas at the lesion site it is markedly increased and tightly wrapped around motoneurons. Together, these findings demonstrate that rMS promotes functional respiratory recovery after cervical SCI through combined enhancement of diaphragmatic motor output and modulation of the inflammatory and extracellular environment. Together, these functional and cellular findings indicate that spinal rMS promotes a permissive, pro-regenerative environment supporting respiratory circuit plasticity. We conclude that rMS significantly enhances ventilatory recovery via reduced inflammatory response and improved intraspinal rewiring after high cervical SCI, suggesting it is a promising non-invasive strategy. The ability of rMS to engage spared respiratory networks and support neuroplasticity highlights its promise as a safe, non-invasive therapeutic strategy with translational potential for rehabilitation of breathing function after SCI.

**One Sentence Summary:** Noninvasive cervical magnetic stimulation improves breathing after spinal cord injury by boosting diaphragm activity and reducing inflammation.

## INTRODUCTION

Spinal cord injury (SCI) affects over 20.6 million people worldwide, with 900,000 new cases reported each year (Ding *et al*, 2022; Qin *et al*, 2024; Safdarian *et al*, 2023). Among these, high cervical SCI often results in chronic respiratory insufficiency and, in many cases, a lifelong inability to breathe independently (Brown *et al*, 2006; Josefson *et al*, 2021; Zimmer *et al*, 2007). Respiratory complications, including pneumonia and bronchitis, are the leading causes of death in both the acute and chronic phases following SCI (Burns, 2007; Johansson *et al*, 2025). Currently, mechanical ventilation remains the standard of care for patients with high-level cervical injury with reduced ventilatory capacity (Korupolu *et al*, 2021; Schreiber *et al*, 2021). While lifesaving, this approach introduces numerous complications, such as frequent suctioning of respiratory secretions, impaired speech and smell, increased risk of infection, and significant psychological and physical burden for both patients and caregivers (Galeiras Vazquez *et al*, 2013; Hendershot & O’Phelan, 2022; Tollefsen & Fondenes, 2012). Moreover, prolonged mechanical ventilation leads to diaphragm disuse and subsequent atrophy, further impeding the chance of spontaneous respiratory recovery (Levine *et al*, 2008; Penuelas *et al*, 2019; Powers *et al*, 2009; Smuder *et al*, 2016).

Diaphragm pacing, which stimulates the phrenic nerve to contract the diaphragm, offers a potential alternative for selected patients. It can reduce ventilator dependence and improve quality of life (Onders *et al*, 2022; Romero-Ganuza *et al*, 2011; Sharma *et al*, 2021). However, this approach requires intact phrenic nerve conduction and a functional lower motor neuron pool. Unfortunately, only about 5% of patients with cervical SCI meet these criteria (Skalsky *et al*, 2015; Vashisht & Chowdhury, 2025), severely limiting its applicability. Consequently, the majority of individuals with high cervical injuries are left without practical options to regain independent breathing.

One of the major barriers to recovery following SCI is the neuroinflammatory response consecutive to the initial injury. The primary injury initiates a cascade of cellular and molecular events that lead to long-lasting secondary damage. Resident microglia become rapidly activated and release proinflammatory cytokines such as TNF-α, IL-1β, and IL-6, aiming at containing tissue damage, promoting debris clearance, but at the same time contributing to demyelination, oxidative stress, and neuronal apoptosis (Hellenbrand *et al*, 2021; Li *et al*, 2022a). Similarly, monocyte-derived macrophages infiltrate the lesion site, sustaining chronic inflammation and interfering with tissue repair. In addition, activated astrocytes undergo hypertrophy and form a dense glial scar, secreting extracellular matrix molecules such as chondroitin sulfate proteoglycans (CSPGs), which are highly repulsive to axonal growth and plasticity (Anderson *et al*, 2016; Silver & Miller, 2004; Wanner *et al*, 2008; Yang *et al*, 2020). These CSPGs, along with other extracellular matrix components, accumulate around neurons to form perineuronal nets (PNNs), a structure that stabilizes synaptic connections but also severely limits synaptic remodeling after injury (Bradbury & Burnside, 2019; Dyck *et al*, 2018; Dyck & Karimi-Abdolrezaee, 2015; Fawcett *et al*, 2022; Sorg *et al*, 2016). Additionally, PDGFRβ-positive pericytes proliferate and contribute to fibrotic scarring, further reinforcing an environment that is non-permissive to regeneration (Birbrair *et al*, 2014; Dias *et al*, 2021; Picoli *et al*, 2019; Yao *et al*, 2022). This chronic neuroinflammatory microenvironment is particularly detrimental to the phrenic motor circuit, which spans the C3 to C5 segments. After cervical SCI, especially in the region around C3/4, the surviving phrenic motor neurons are surrounded by reactive glia, inflammatory mediators, and dense PNNs that limit their ability to reconnect or reorganize (Lukacova *et al*, 2021; Sánchez-Ventura *et al*, 2023; Windelborn & Mitchell, 2012). These cellular and molecular events act synergistically to prevent meaningful respiratory recovery. Thus, strategies that can modulate inflammation and reduce extracellular matrix barriers are critical for functional repair.

Currently, neurostimulation emerges as a promising therapeutic strategy for enhancing neural plasticity after SCI (Dorrian *et al*, 2023; Inanici *et al*, 2018; Van Steenbergen *et al*, 2023). Among available techniques, repetitive magnetic stimulation (rMS) is particularly attractive due to its non-invasive nature and established safety in clinical settings (Mann & Malhi, 2025). Originally developed as a treatment for depression (Loo *et al*, 2008), rMS modulates cortical excitability and synaptic plasticity, and has been approved for clinical use by regulatory agencies worldwide (Cotovio *et al*, 2023; Lenz *et al*, 2016). In animal models, repetitive trans-spinal magnetic stimulation (rTSMS) has shown the capacity to enhance functional recovery after SCI, reduce glial reactivity, and stimulate axonal sprouting (Jiang *et al*, 2024; Liu *et al*, 2020; Robac *et al*, 2021). Several studies have also reported that rMS can downregulate inflammatory signaling and reduce microglial activation in models of traumatic brain injury and SCI (O’Leary *et al*, 2025; Sasso *et al*, 2016; Sekar *et al*, 2019; Toledo *et al*, 2021; Zong *et al*, 2020).

Importantly, in cervical SCI models, rMS applied to the spinal cord has been shown to increase diaphragm activity, likely by enhancing excitability of the phrenic circuit and possibly through modulation of the local microenvironment (Lee & Vinit, 2024; Lv *et al*, 2023; Michel-Flutot *et al*, 2021). Despite these encouraging results, rMS protocols vary widely across studies, including differences in stimulation frequency, intensity, duration, and anatomical targeting (Dufor *et al*, 2023; Kim *et al*, 2020). Such variability complicates comparisons between studies and limits mechanistic interpretation, particularly in the context of respiratory recovery. Stimulation frequency is a critical parameter, as it strongly influences the direction and magnitude of plastic changes (Brihmat *et al*, 2022; Torii *et al*, 2012). High-frequency protocols, typically at or above 10 Hz, are generally associated with increase neuronal excitability and modulate the balance between excitatory and inhibitory inputs, for example by enhancing phrenic motor neuron output and attenuating GABAergic inhibition (Lenz *et al*., 2016; Michel-Flutot *et al*., 2021), whereas lower frequencies tend to produce inhibitory outcomes.

In the present study, we therefore selected a 10 Hz rMS protocol, as this frequency has been consistently shown to enhance excitability and promote plasticity in both cortical and spinal circuits. Importantly, 10 Hz stimulation has also been used safely in clinical and preclinical settings and has demonstrated efficacy in modulating motor and respiratory-related pathways.

We sought to determine whether 10 Hz rMS could reduce neuroinflammation and promote respiratory recovery in a mouse model of C3/4HC. We hypothesized that chronic cervical rMS would attenuate microglial and astrocyte activation, decrease CSPG deposition and/or PNN formation, leading to a more preserved phrenic motor network. Furthermore, we aimed to evaluate whether these cellular changes translate into improved diaphragm function. By targeting both the inflammatory and functional components of injury, this study explores the potential of rMS as a non-invasive therapeutic strategy to restore breathing in patients with high cervical SCI.

## RESULTS

### Physiological Effect and Histological Analysis in C3HC Mice Treated with rMS or Sham Stimulation

A transient decrease in body weight was observed in both C3HC groups (sham rMS and rMS) during the first week following surgery which then re-increased progressively similarly between groups. By 21 days post-injury, animals had reached levels comparable to their preoperative baseline. No significant difference in body weight was found between the sham rMS and rMS groups at any of the three time points (D0, D7, or D21; Fig. 1C, Supplementary Table 1). Histological quantification of the lesion extent revealed no significant difference between groups, with comparable percentages of injured hemicord in the sham rMS and rMS-treated animals (79.0 ± 6.5% vs. 74.8 ± 8.3%).

**Fig. 1.**
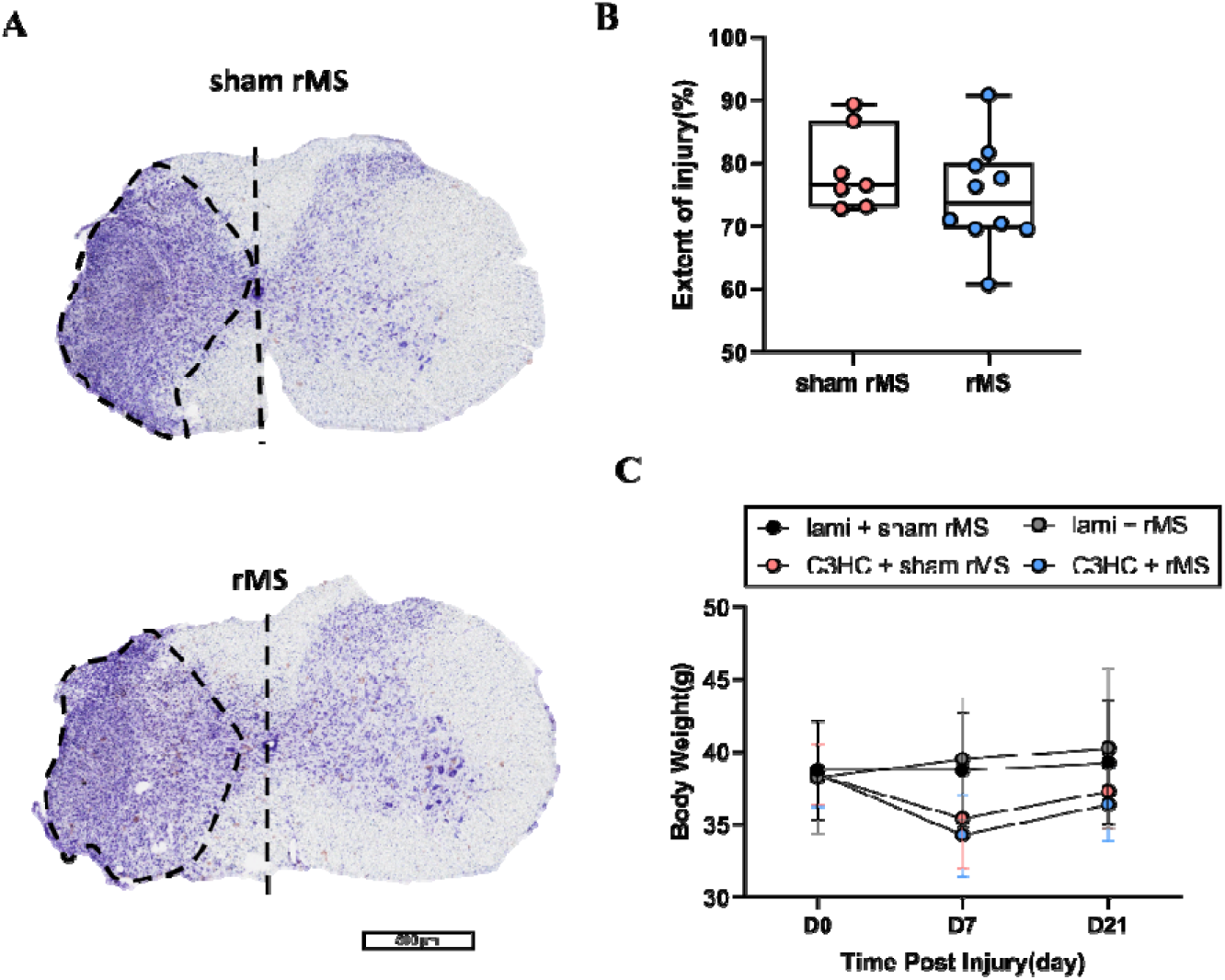
Examples of extent of injury following a C3 hemicontusion treated with rMS or sham stimulation. **(A)** Representative examples of cresyl violet staining at the injured site for in sham rMS (top) and rMS treated groups (bottom). The lesion area is delineated by the dotted black outline; the vertical dashed line indicates the spinal midline. Scale Bar: 500 μm. **(B)** Quantification of injury extent, calculated as the area of the lesion relative to the total area of the ipsilateral hemi-spinal cord (set as 100%). Statistical analysis was performed using the unpaired Student’s t-test. **(C)** Body weight measurements of animals across the four experimental groups including lami + sham rMS, lami + rMS, C3HC + sham rMS, C3HC + rMS group at three time points: baseline (D0), 7 days post-injury (D7), and 21 days post-treatment (D21). Data represent mean ± SD. Statistical analysis was performed using the Two way repeated-measures (RM) ANOVA followed by Tukey’s post hoc multiple comparisons test.

### Effects of rMS on Respiratory Function

Plethysmography measurements performed under light anesthesia (1% isoflurane) are presented in Fig. 2 and table S1. In the whole group of mice, cervical SCI induced a significant reduction in tidal volume (VT) in all animals when corrected for body weight. By day 7 (D7) post-injury, VT had decreased by approximately 44% in the Sham group (5.81 ± 0.88 vs. 3.08 ± 0.88 µL/g, *p*<0.01), and by 32% in the rMS group (5.71 ± 0.61 to 3.88 ± 1.01µL/g, *p*<0.001), with no significant difference between groups. However, recovery profiles differed by day 21 (D21). VT was significantly higher in the rMS group (rMS vs sham rMS: 6.02 ± 1.15 vs. 3.87 ± 1.01 µL/g, *p*<0.01) compared to the sham group with limited spontaneous recovery. There was no significant change observed for Breathing frequency (Bf) between groups or time points. As a result, similar changes were observed for Minute ventilation (VE). Mice treated rMS group recovered faster than those in the Sham group, and a significant difference was observed between groups (rMS vs sham rMS: 36.53 ± 11.56 vs. 21.94 ± 6.48 mL/min, p<0.05).

**Fig. 2.**
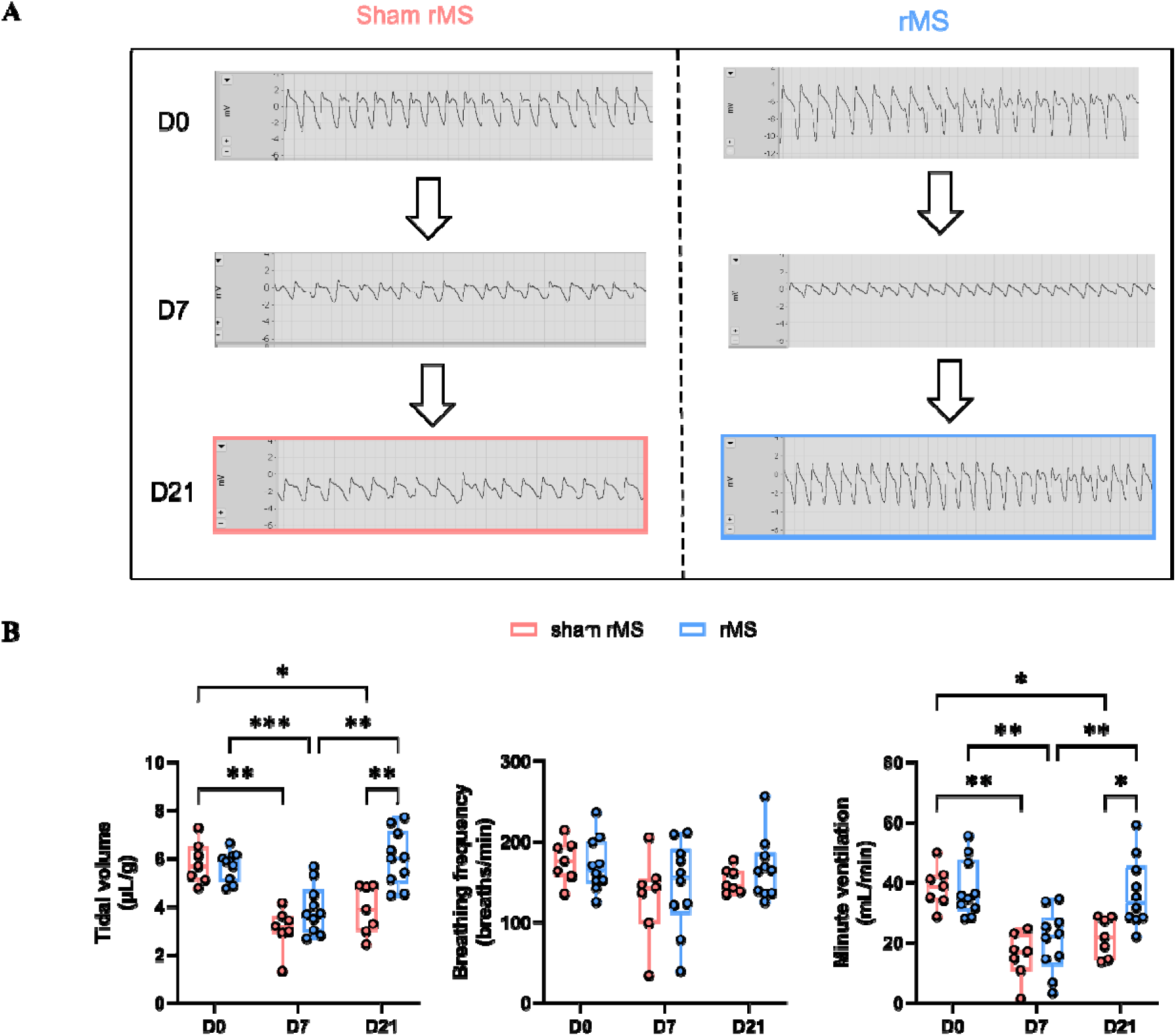
Respiratory parameters in C3HC mice treated with rMS or sham stimulation under light anesthesia (1% isoflurane). Histograms (Min to max, Box and Whiskers) present tidal volume (VT), breathing frequency (Bf) and minute ventilation (VE) measured before surgery (D0), 7 days after injury (D7), and 21 days after injury (D21) in C3HC mice treated with sham rMS or rMS. Statistical analysis was performed using the Two way repeated-measures (RM) ANOVA followed by Tukey’s post hoc multiple comparisons test. *** *p* < 0.001, ** *p* < 0.01, * *p* < 0.05.

During hypercapnic challenge with 5% CO (Fig. 3), VT did not differ significantly between baseline and 7 days post-injury in either the sham (8.05 ± 1.63 vs. 5.92 ± 2.29 µL/g, n.s) or rMS groups (8.26 ± 2.19 vs. 6.12 ± 3.13 µL/g, n.s). In the rMS group, VT significantly increased by day 21 compared to day 7 (p < 0.01), while no significant recovery was observed in the sham group (rMS vs sham rMS: 10.09 ± 3.71 vs. 7.36 ± 2.52 µL/g, n.s). Although VT tended to be higher in the rMS group at day 21, the between-group difference was not statistically significant. Bf remained unchanged across time points and treatment groups. VE followed a similar pattern to VT, with a significant reduction at D7 in rMS groups (48.44 ± 13.20 vs. 31.05 ± 19.45 mL/min, p < 0.05) and with a similar trend in sham rMS group (49.01 ± 13.40 vs. 31.52 ± 16.62 mL/min, p = 0.08). VE significantly increased in the rMS group at D21 compared to D7 (p < 0.001), while sham mice showed no recovery. No significant difference in VE was found between groups at D21, although a trend toward higher values was observed in the rMS group (rMS vs sham rMS: 54.58 ± 23.82 vs. 40.02 ± 14.72 mL/min, n.s).

**Fig. 3.**
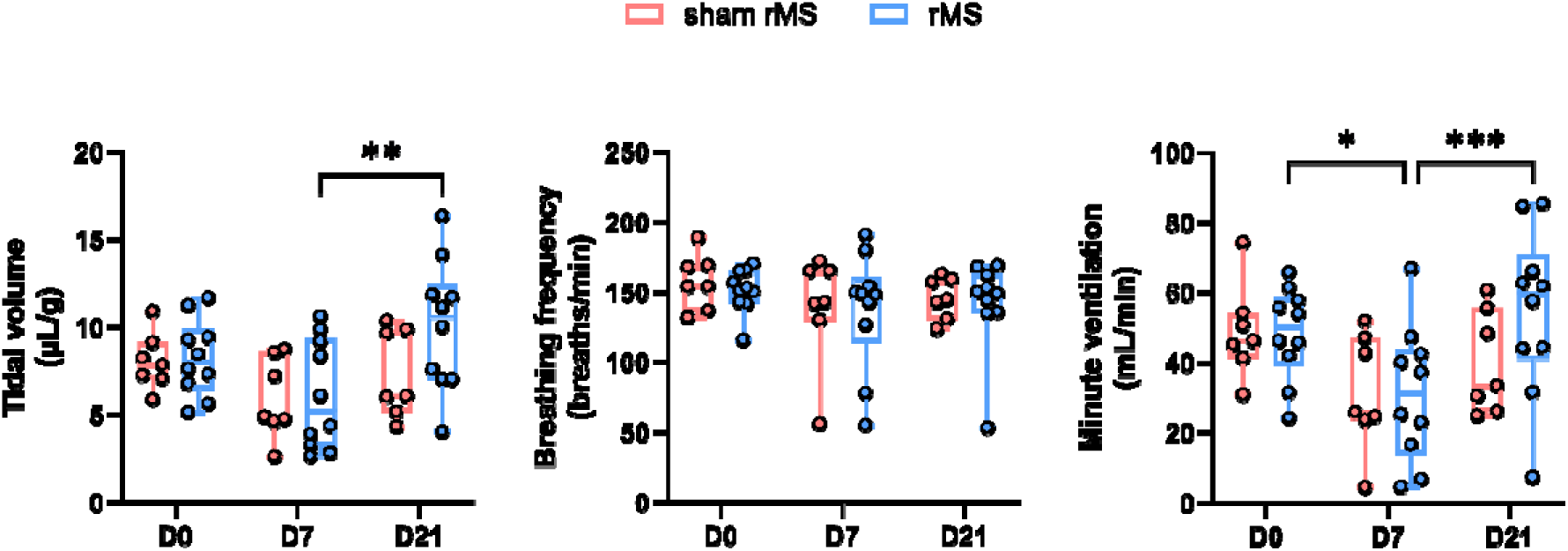
Respiratory parameters in C3HC mice treated with rMS or sham stimulation in the 5% CO□ phase. Histograms (Min to max, Box and Whiskers) present tidal volume (VT), breathing frequency (Bf) and minute ventilation (VE) measured before surgery (D0), 7 days after injury (D7), and 21 days after injury (D21) in C3HC mice treated with sham rMS or rMS. Statistical analysis was performed using the Two way repeated-measures (RM) ANOVA followed by Tukey’s post hoc multiple comparisons test. *** p < 0.001, ** p < 0.01, * p < 0.05.

Under light anesthesia (1% isoflurane) after hypercapnic challenge (Fig. 4), VT tended to decrease from baseline to day 7 in both sham (4.57 ± 1.08 vs. 3.03 ± 0.58 µL/g, p = 0.05) and rMS groups (4.48 ± 1.23 vs. 3.27 ± 1.38 µL/g, n.s). By day 21, VT in the rMS group showed a trend toward recovery, with higher values compared to the sham group, although the difference was not statistically significant (rMS vs sham rMS: 4.31 ± 1.31 vs. 3.58 ± 1.12 µL/g, n.s). Bf remained stable across timepoints without intergroup differences. VE also declined at D7 in both sham rMS (26.65 ± 7.01 vs. 14.91± 5.69 mL/min, p = 0.08) and rMS (25.60 ± 11.05 vs. 14.48± 7.00 mL/min, p = 0.05) groups. At D21, VE showed partial recovery in the both groups (rMS vs sham rMS: 18.51 ± 6.36 vs. 18.44 ± 5.15 mL/min, n.s). Although the group difference was not significant, the upward trend in VE paralleled the improvement observed in VT.

**Fig. 4.**
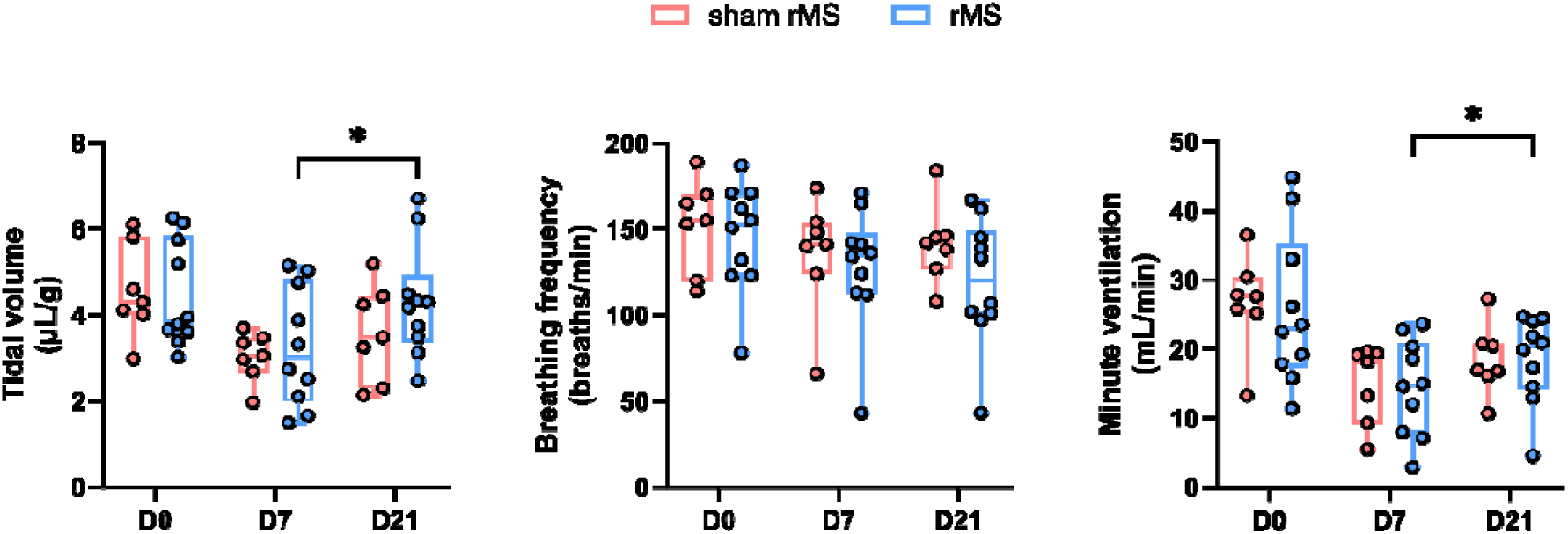
Respiratory parameters in C3HC mice treated with rMS or sham stimulation in the 1% isoflurane (after hypercapnic challenge). Histograms (Min to max, Box and Whiskers) present tidal volume (VT), breathing frequency (Bf) and minute ventilation (VE) measured before surgery (D0), 7 days after injury (D7), and 21 days after injury (D21) in C3HC mice treated with sham rMS or rMS. Statistical analysis was performed using the Two way repeated-measures (RM) ANOVA followed by Tukey’s post hoc multiple comparisons test. * p < 0.05.

During spontaneous breathing in ambient air (normal poikilocapnic breathing) (Fig. 5), respiratory parameters remained relatively stable in the sham rMS group throughout the post-injury period. In contrast, animals treated with rMS showed significant recovery of VT and VE by day 21 compared to baseline (D0 vs D21: p < 0.05), indicating a progressive improvement in respiratory capacity. Although no significant difference was observed between rMS and sham groups at each individual time point, the within-group recovery trend in the rMS group suggests beneficial effect of rMS on restoring respiratory output under physiological breathing conditions. Breathing frequency (Bf) significantly change after day 7 and day 21 in sham rMS group (D0 vs D7 vs D21: 175 ± 29 vs. 210 ± 17 vs. 222 ± 30 breaths/min, p < 0.05), whereas a nonsignificant upward trend in rMS group (D0 vs D7 vs D21: 203 ± 36 vs. 229 ± 18 vs. 230 ± 31 breaths/min, n.s).

**Fig. 5.**
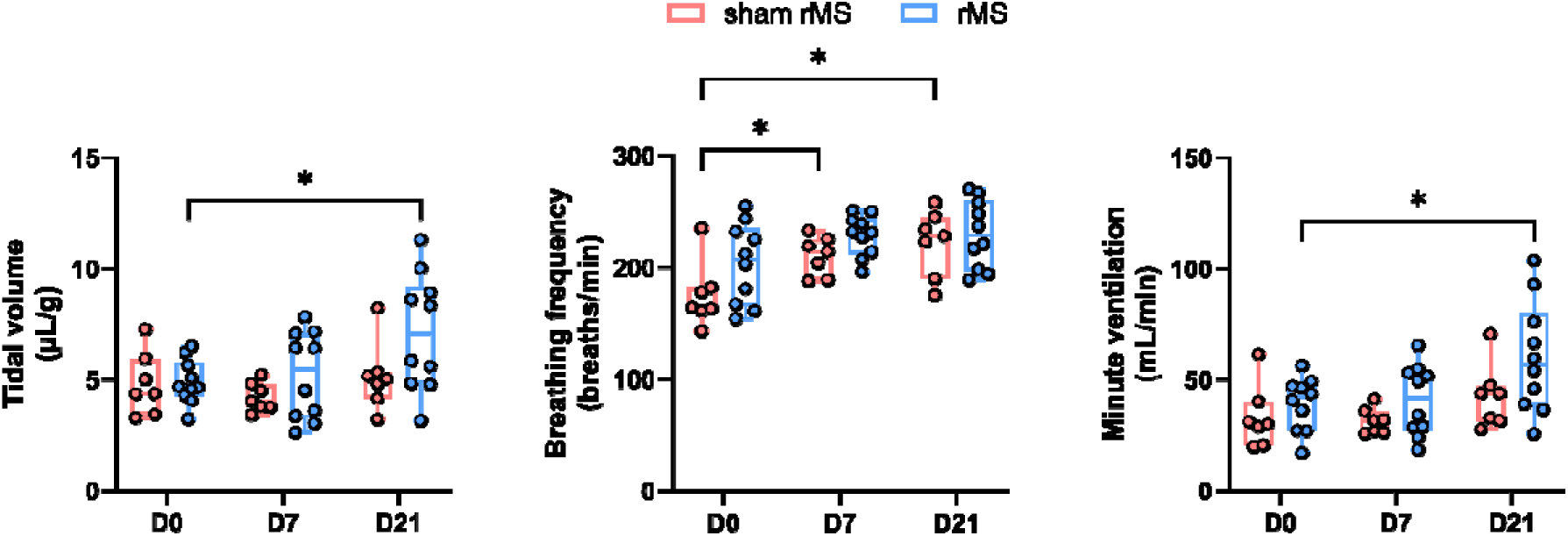
Respiratory parameters in C3HC mice treated with rMS or sham stimulation during spontaneous (poikilocapnic) breathing. Histograms (Min to max, Box and Whiskers) present tidal volume (VT), breathing frequency (Bf) and minute ventilation (VE) measured before surgery (D0), 7 days after injury (D7), and 21 days after injury (D21) in C3HC mice treated with sham rMS or rMS. Statistical analysis was performed using the Two way repeated-measures (RM) ANOVA followed by Tukey’s post hoc multiple comparisons test. * p < 0.05.

### Effects of rMS on Diaphragm Activity

To investigate the mechanistic basis of improved respiratory function, we assessed dia-EMG activity across three anatomical regions (ventral, medial, and dorsal) for both intact and injured sides (Fig. 6). On the injured side, rMS significantly preserved dia-EMG amplitude close to control values, while it tended to decrease in the sham-rMS group after C3HC injury. In the ventral region of the diaphragm, a reduction in dia-EMG activity was observed in the C3HC + sham rMS group compared to the lami + sham rMS group (0.40 ± 0.27 vs. 0.77 ± 0.38 μV·s·s). Remarkably, rMS treatment effectively preserved dia-EMG activity, restoring it to 0.77 ± 0.33 μV·s·s, a level significantly higher than the C3HC + sham-rMS injury group (p < 0.05). In the medial region, a similar pattern was observed, with reduced activity in the C3HC + sham rMS group (0.47 ± 0.3 vs. 0.68 ± 0.4 μV·s·s) compared to the lami + sham rMS group and restored following rMS treatment (0.86 ± 0.41 μV·s·s; p < 0.05). No significant differences were observed in the dorsal region among groups.

**Fig. 6.**
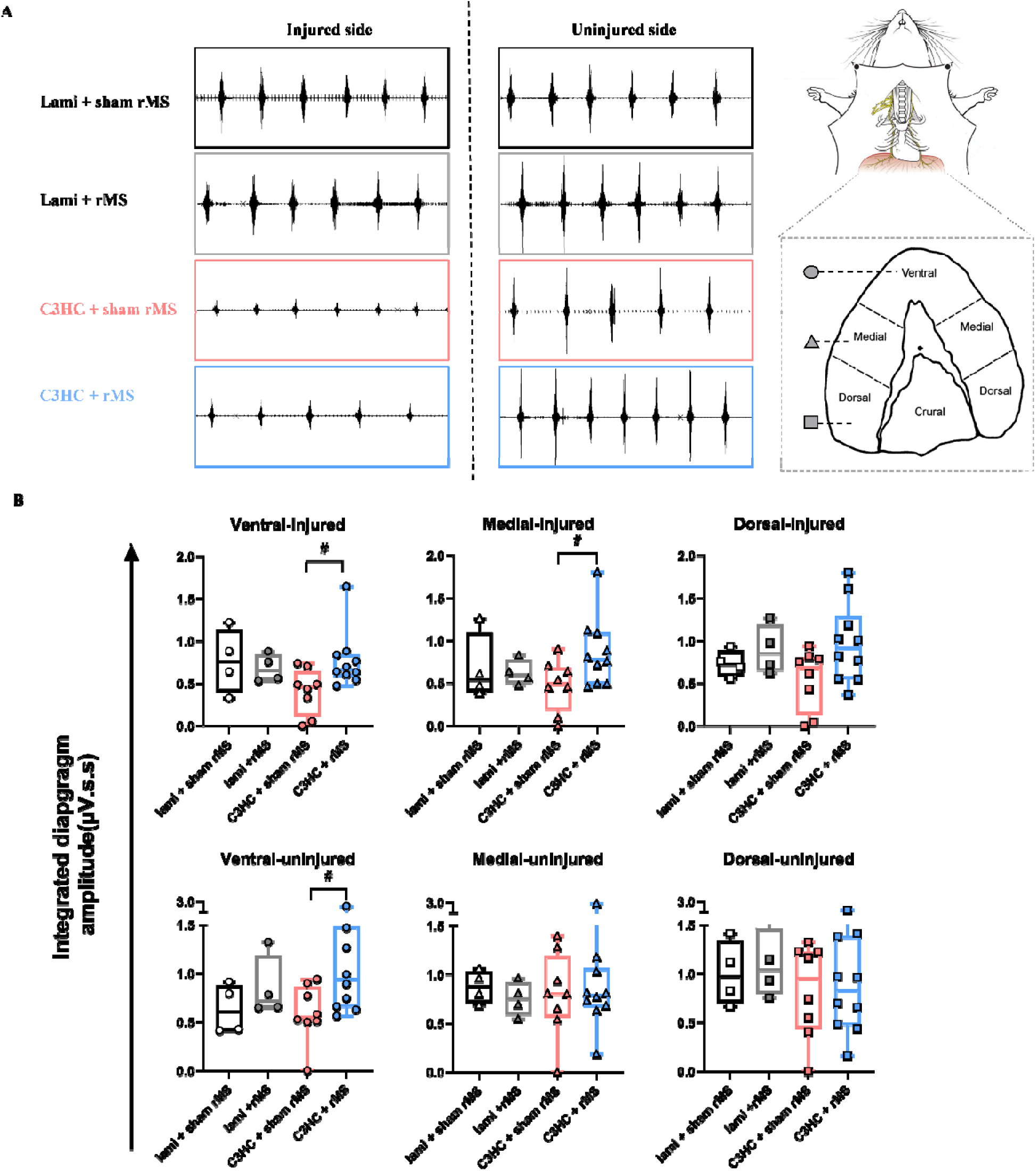
Diaphragm activity in lami or C3HC mice treated with rMS or sham stimulation. **(A)** Representative examples of diaphragmatic EMG in mice with sham rMS and rMS treated groups from the ventral part of diaphragm on the injured side. **(B)** Quantification of peak amplitude of double integrated diaphragm EMG (expressed in μV·s·s) on both the injured and uninjured sides. All values have been corrected for the amplifier gain factor to represent biological source voltage. Data are presented as Histograms (Min to max, Box and Whiskers) in four experimental groups: lami + sham rMS (black), lami + rMS (gray), C3HC + sham rMS (red), and C3HC + rMS (blue). Each dot represents one animal. Comparisons are shown across the ventral, medial, and dorsal regions of the diaphragm. Statistical significance between the C3HC + sham rMS and C3HC + rMS groups was assessed using an unpaired student’s t test(^#^), ^#^ *p* < 0.05.

On the uninjured side, rMS also enhanced diaphragm activity in the ventral region. Dia-EMG amplitude significantly increase in the C3HC + rMS group compared to the C3HC + sham rMS group (1.1 ± 0.57 vs. 0.59 ± 0.3 μV·s·s), while the laminectomy group showed a baseline level (0.85 ± 0.32 vs. 0.64 ± 0.26 μV·s·s) and without significant difference. This increase in activity may reflect a compensatory enhancement in the contralateral diaphragm. No significant differences were observed among groups in the middle and dorsal regions on the uninjured side.

### rMS Modulates Scar-associated and Inflammatory Cellular Responses After C3HC

#### rMS Reduces Fibrotic Cells and Astrocytic Scar Formation

Immunohistofluorescence analysis of PDGFRβ and GFAP at the lesion level is shown in Fig. 7, and around lesion level are presented in fig. S1. PDGFRβ and GFAP were co-expressed within the fibrotic and astroglial components of the glial scar. A significantly higher percentage area of PDGFRβ-positive staining was observed in the C3HC + sham rMS group compared to the laminectomy group (p < 0.01). rMS treatment markedly reduced the PDGFRβ-positive area, with a significant difference between the sham and rMS-treated C3HC groups (20.98 ± 2.98 % vs. 12.89 ± 2.58 %, p < 0.001). Similarly, PDGFRβ-positive cells counts were elevated in the C3HC + sham rMS group compared to the laminectomy group and significantly reduced following rMS treatment (278 ± 56 vs. 216 ± 47 cells, p < 0.05). Furthermore, GFAP-positive staining was significantly increased in the C3HC injury group compared to the laminectomy group (p < 0.05). The GFAP-positive area was higher in the C3HC + sham rMS group and tended to be decrease by rMS treatment (51.57 ± 14.78 % vs. 40.70 ± 9.80 %, p = 0.08). Consistently, the number of GFAP-positive cells was significantly higher at the lesion site in the sham group and was reduced by rMS (584 ± 80 vs. 469 ± 90 cells, p < 0.05). Concordant reductions around the lesion level are shown in fig. S1.

**Fig. 7.**
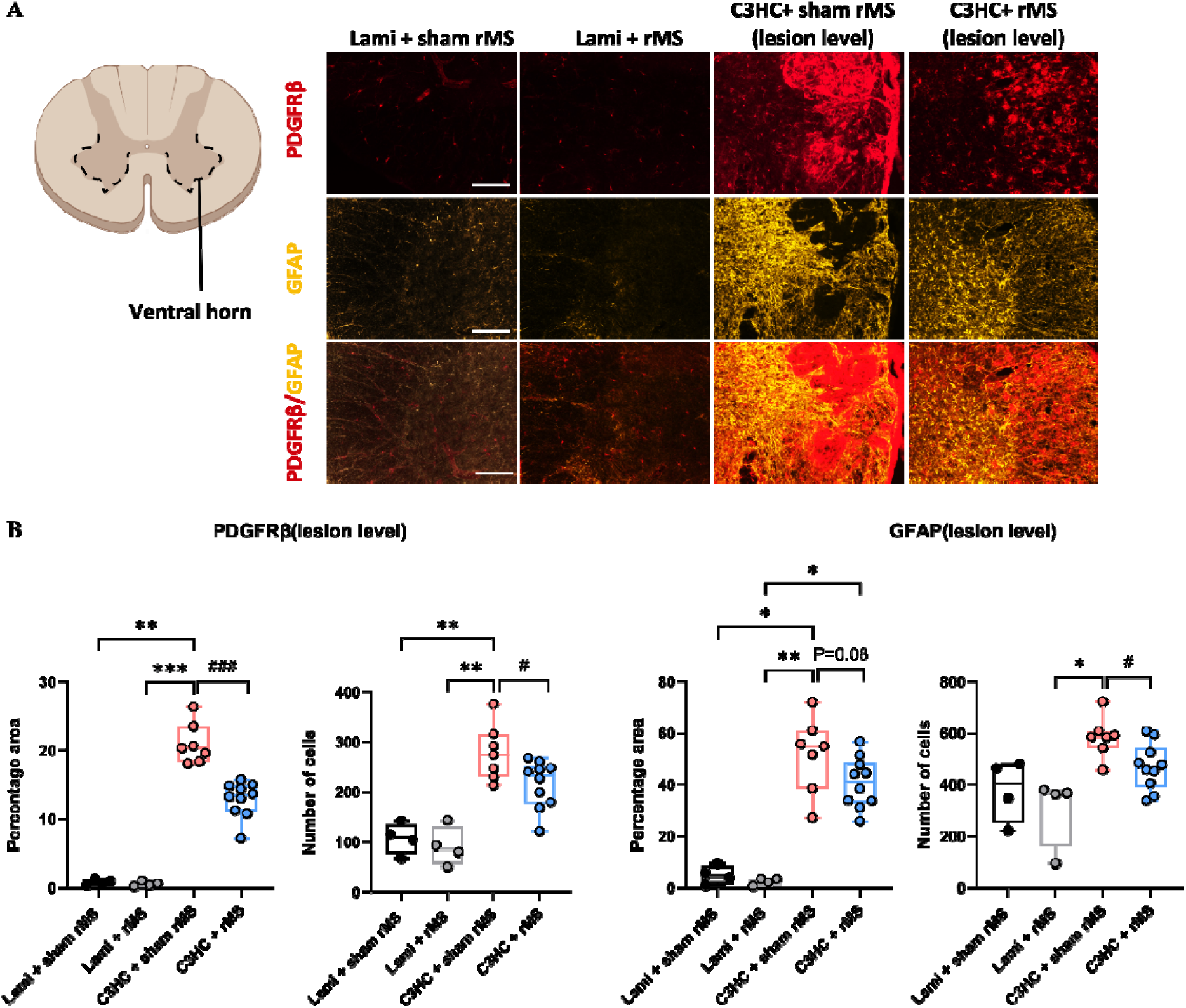
PDGFRβ and GFAP immunoreactivity at 21 days post-injury in the cervical spinal cord lesion level. **(A)** Representative immunofluorescence images showing PDGFRβ (red), GFAP (yellow), and merged channels in the lesion area of Laminectomy + sham rMS, Laminectomy + rMS, C3HC + Sham rMS, and C3HC + rMS-treated animals at 21 days post-injury. (Scale bar: 500μm) **(B)** Quantification of PDGFRβ+ percentage area, GFAP+ percentage area, and cell number in the lesion site at the ventral part of spinal cord. Each dot represents one animal. Histograms (Min to max, Box and Whiskers) show data distribution across the four groups. Statistical significance across the four groups was determined using a Kruskal-Wallis test followed by Dunn’s post-hoc multiple comparisons test (*) *p<0.05, **p<0.01, ***p<0.001, while direct comparisons between the C3HC + sham rMS and C3HC + rMS-treated groups were specifically performed using a Mann-Whitney test (^#^) ^#^p<0.05, ^##^p<0.01, ^###^p<0.001.

#### rMS Attenuates Microglial/Macrophage Activation

CD68 and Iba1, both associated with activated pro-inflammatory microglia/macrophages, were used to assess neuroinflammation (Fig. 8). At the lesion level, the CD68-positive area was significantly increased in the C3HC + sham rMS group compared to the laminectomy group (p < 0.01). rMS treatment significantly reduced the percentage area of CD68-positive labeling in the C3HC model (21.22 ± 5.04 % vs. 14.94 ± 3.92%, p < 0.05). The number of CD68-positive cells decreased following rMS, (423 ± 117 vs. 304 ± 55 cells, p = 0.05). The percentage area of Iba1-positive staining was significantly higher in the C3HC + sham rMS group compared to the rMS-treated group (34.09 ± 9.15 % vs. 24.70 ± 6.53 %, p < 0.01). The number of Iba1-positive cells was also likewise reduced by rMS treatment (417 ± 88 vs. 329 ± 74 cells, p = 0.05). Similar trends were observed around lesion level (Supplementary Figure S2). However, for CD206, a marker of anti-inflammatory microglia/macrophages, the percentage area and the number of CD206-positive cells did not differ between groups (1.21 ± 1.64 % vs. 3.51 ± 6.26 %; 65 ± 77 vs. 66 ± 76 cells). A reduction in fibrotic and glial scarring, as well as pro-inflammatory microglial activation, was also observed around the lesion site following rMS (Supplementary Figure S4).

**Fig. 8.**
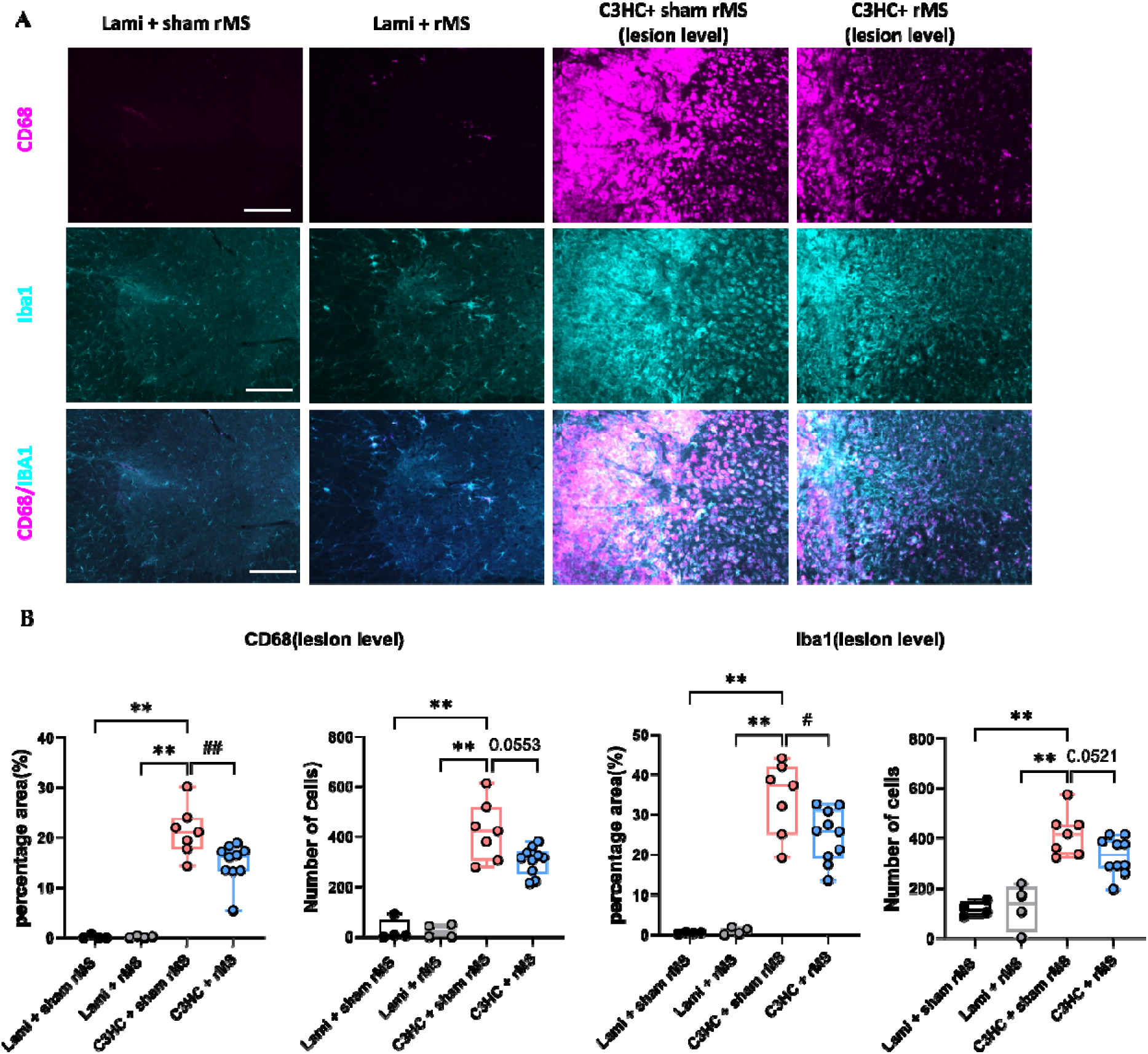
CD68 and Iba1 immunoreactivity at 21 days post-injury in the cervical spinal cord lesion level. **(A)** Representative immunofluorescence images showing CD68 (purple), Iba1 (cyan), and merged channels in the lesion area of Laminectomy + sham rMS, Laminectomy + rMS, C3HC + Sham rMS, and C3HC + rMS-treated animals at 21 days post-injury. Scale bar: 500 µm. **(B)** Quantification of CD68+ percentage area, Iba1+ percentage area, and cell number in the lesion site at the ventral part of spinal cord. Each dot represents one animal. Histograms (Min to max, Box and Whiskers) show data distribution across the four groups. Statistical significance across the four experimental groups was determined using a Kruskal-Wallis test followed by Dunn’s post-hoc multiple comparisons test (*) *p<0.05, **p<0.01, ***p<0.001, while direct comparisons between the C3HC + sham rMS and C3HC + rMS-treated groups were specifically performed using a Mann-Whitney test (^#^) ^#^p<0.05, ^##^p<0.01, ^###^p<0.001.

**Fig. 9.**
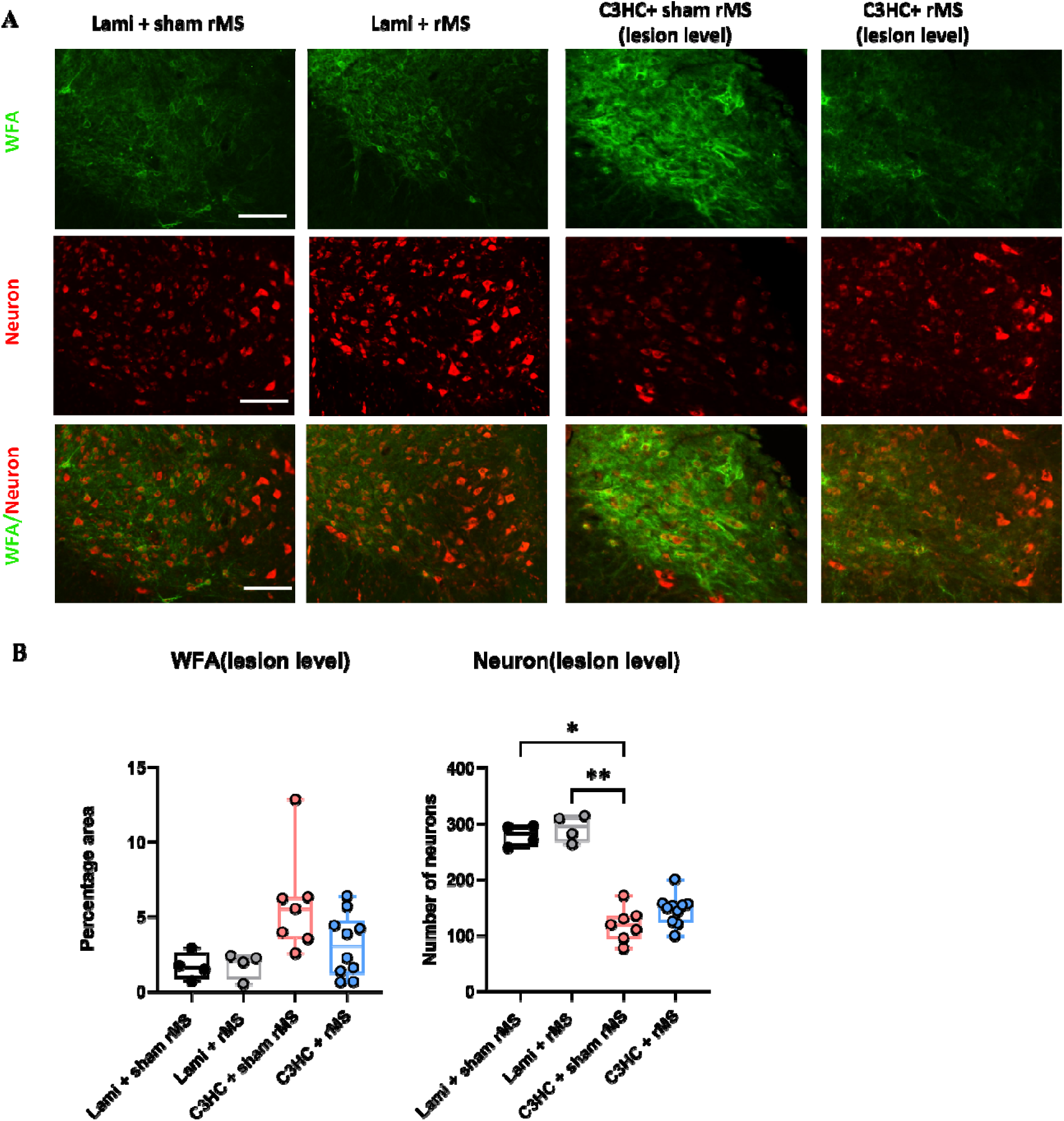
WFA staining and ventral neuron survival at 21 days post-injury in the cervical spinal cord lesion level. **(A)** Representative immunofluorescence images showing WFA (green), NeuroTrace™ 435/455 (neuronal marker, red), and merged channels in the lesion area of Laminectomy + sham rMS, Laminectomy + rMS, C3HC + sham rMS, and C3HC + rMS-treated animals at 21 days post-injury. NeuroTrace™ 435/455, which possesses intrinsic blue fluorescence, has been pseudocolored red using image processing software to enhance visualization and contrast within the merged channels. Scale bar: 500 µm. **(B)** Quantification of WFA-positive percentage area, number of ventral neurons at the lesion site. Each dot represents one animal. Histograms (Min to max, Box and Whiskers) show data distribution across the four groups. Statistical significance across the four experimental groups was determined using a Kruskal-Wallis test followed by Dunn’s post-hoc multiple comparisons test (*) *p<0.05, **p<0.01.

**Fig. 10.**
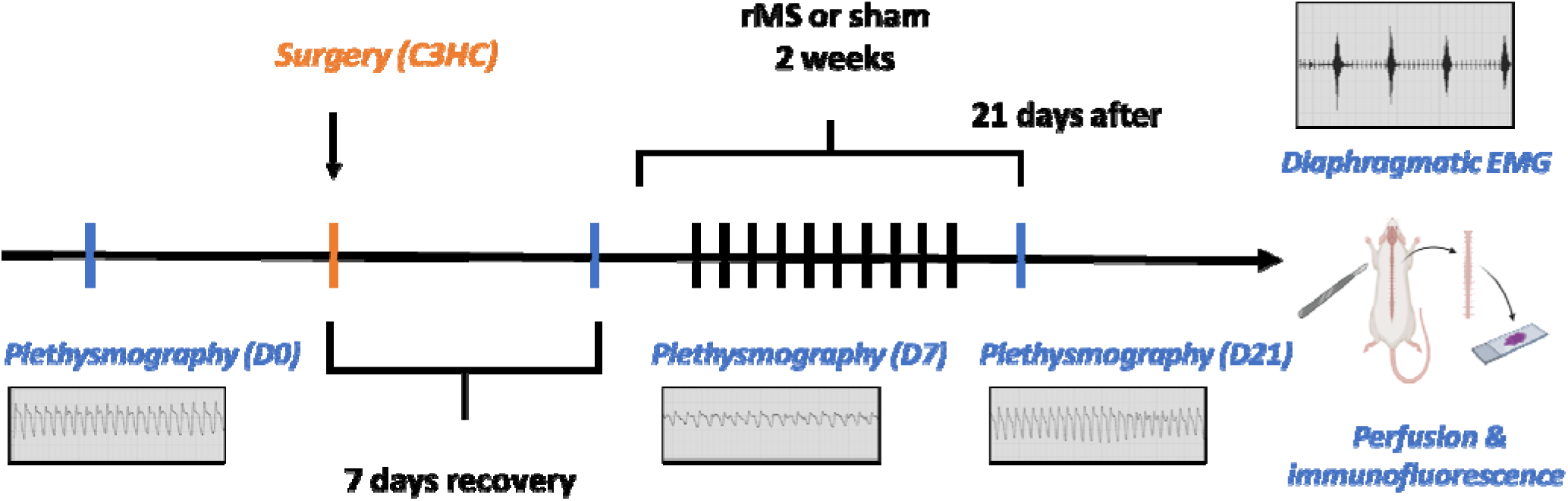
Experimental timeline. Schematic representation of the experimental design to evaluate the effects of rMS on respiratory function after cervical spinal cord injury. Baseline respiratory function was performed before surgery (Day 0), followed by C3/4 hemi-contusion or laminectomy (Day 1). Respiratory function was reassessed on Day 7. rMS or sham stimulation was applied daily from Day 8 to Day 21. Final assessments were performed, including respiratory function, diaphragm activity, and tissue collection for immunofluorescence.

**Figure 11.**
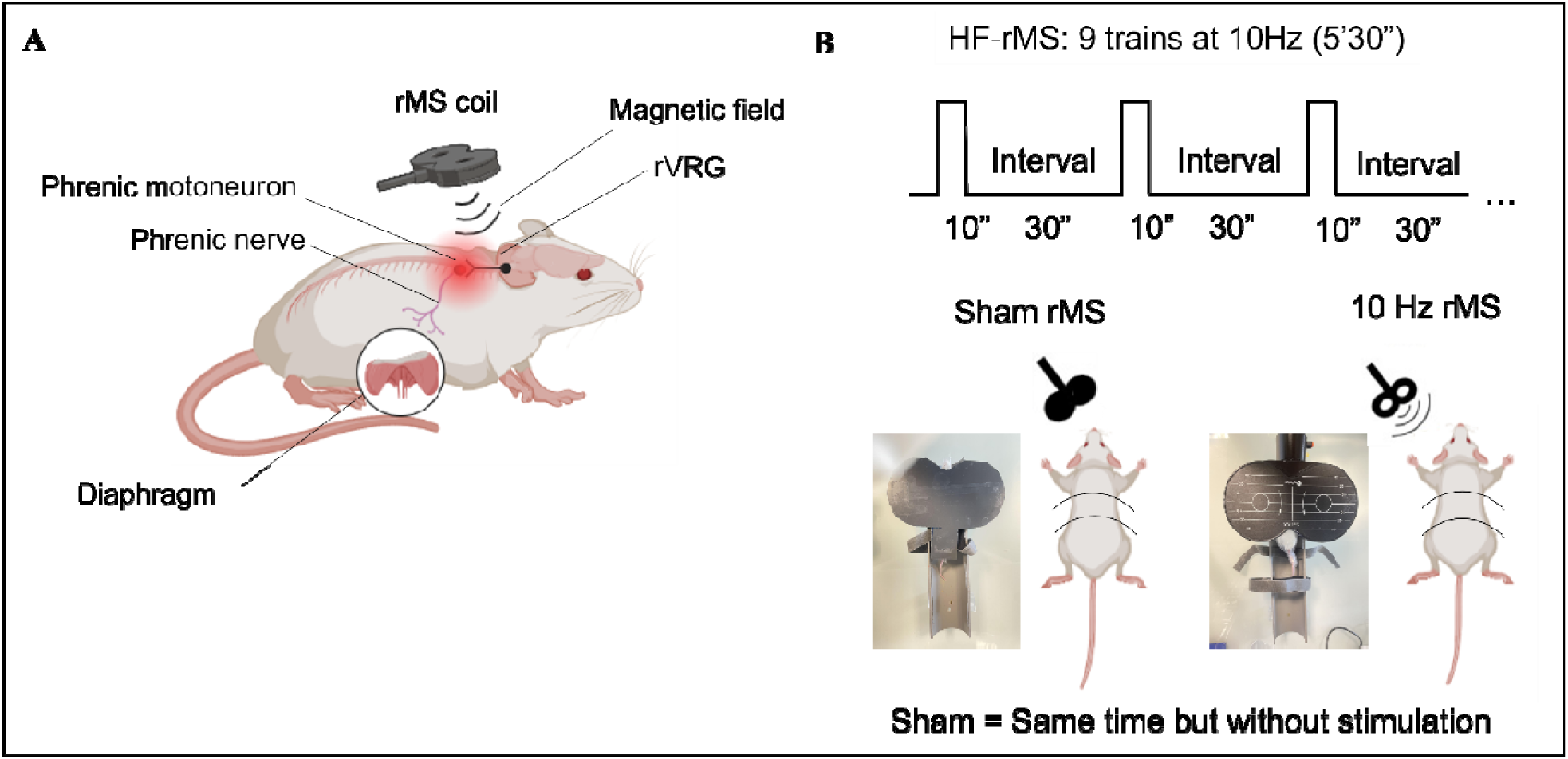
rMS experimental setup and stimulation protocol. **(A)** Schematic of the phrenic motor circuit and rMS coil placement targeting the rVRG and cervical phrenic motoneurons. The diaphragm is shown as the functional output. **(B)** HF-rMS protocol parameters (top) and experimental configuration for sham and active 10 Hz stimulation in awake, restrained mice (bottom).

#### rMS Modulates WFA labeled Perineuronal Nets and Neuronal Preservation

WFA staining showed a trend toward increased CSPG-rich perineuronal net (PNN) signal in the C3HC + sham rMS group compared to the lami + sham rMS (5.86 ± 3.39 % vs. 1.71 ± 0.91%), with a trend toward reduction following rMS treatment (5.86 ± 3.39 % vs. 3.10 ± 2.09 %, p =0.1). In parallel, C3HC injury led to a clear reduction in neuronal density (280 ± 19 vs. 119 ± 30 cells, p < 0.05) compared to the Lami + sham rMS group, and neurons in the ventral horn tended to be higher at the lesion site in the rMS group compared to sham stimulation group (119 ± 30 vs. 146 ± 27 cells, p = 0.08). Morphologically, WFA-labeled CSPGs appeared forming dense, cage-like structures condensed around the somata of large ventral horn neurons in sham-stimulated injury animals. In contrast, rMS treatment showed a trend toward reducing this peri-neuronal condensation. Similar trends were observed around lesion level without significance (fig. S3).

## DISCUSSION

Our study shows for the first time that rMS can improve tidal volume recovery after cervical SCI. Indeed, spontaneous recovery remained limited without treatment, whereas rMS accelerated and amplified functional restoration. These functional improvements occurred alongside reduced glial and fibrotic scarring as well as suppression of chronic M1-like activation, suggesting that rMS mitigates inhibitory and cytotoxic cues that normally consolidate lesion pathology. Moreover, rMS showed a trend toward reducing perineuronal net density and was associated with a trend toward neuronal survival, which suggests that rMS may stabilize spared phrenic motoneurons and may facilitate the re-establishment of functional connections within the phrenic motor pool. These effects, likely arising from the modulation of the neuroinflammatory environment, provide a plausible mechanism by which rMS supports the restoration of respiratory capacity.

### Neuroprotection via reduced glial/fibrotic scarring

At 21 days post-injury, the spinal inflammatory response is transitioning from the subacute to the early chronic phase, a stage where the glial and fibrotic scar consolidates into a persistent barrier to repair (Hellenbrand *et al*., 2021; Orr & Gensel, 2018). This scar, formed by proliferating PDGFRβ+ fibroblasts and GFAP+ astrocytes, is known to constrain axonal sprouting and limit the reconnection of spared neural circuits. (Bradbury & Burnside, 2019; Li *et al*, 2022b; Orr & Gensel, 2018; Pang *et al*, 2021). In this study, rMS appeared to shift this balance toward a less restrictive milieu by reducing both PDGFRβ+ fibroblasts and GFAP+ labeled cells. Rather than simply being an anatomical structure, the scar functions as a dynamic cellular and extracellular interface that either fosters or obstructs regeneration depending on its composition. Intriguingly, other studies using trans-spinal magnetic stimulation have reported inhibition of fibrotic scar by increasing in turn astrogliosis reactivity (Chalfouh *et al*, 2020; Robac *et al*., 2021). However, astrocytes are highly heterogeneous in their responses after SCI (Qian *et al*, 2025). On one hand, reactive astrocytes rapidly proliferate and upregulate GFAP to form the glial scar, which physically and chemically restricts axonal growth (Perez-Gianmarco & Kukley, 2023). On the other hand, a subset of astrocytes can adopt a protective role, limiting lesion expansion, isolating inflammatory infiltrates, and providing trophic support (Faulkner *et al*, 2004). Thus, an apparent increase in astrogliosis in some contexts may actually reflect the predominance of this reparative, neuroprotective phenotype (Wang *et al*, 2022). In contrast, the reduced GFAP signal observed in our study may indicate that rMS specifically dampened the maladaptive component of astrocytic activation that contributes to deposition of inhibitory extracellular matrix molecules such as chondroitin sulfate proteoglycans (CSPGs), which are known to inhibit axonal growth and synaptic remodeling since we also observed a reduction in CSPG (Francos-Quijorna *et al*, 2022; Yang *et al*, 2024).

### Suppression of pro-inflammatory microglia/macrophage activation

Pro-inflammatory Microglia and Macrophages are known to remain persistently activated after SCI within the lesion environment, contributing to the maintenance of glial and fibrotic scarring (Bradbury & Burnside, 2019; Pang *et al*., 2021). In the present study, rMS significantly suppressed this chronic inflammatory activation, as reflected by lower CD68+ and Iba1+ labeling, suggesting a shift away from the deleterious M1-like phenotype. These findings are consistent with prior reports that repetitive transcranial magnetic stimulation (rTMS) downregulates microglial activity and suppresses key pro-inflammatory cytokines such as IL-1β, TNF-α, and IL-6 (Guo *et al*, 2023), while promoting anti-inflammatory mediators including IL-10 and neurotrophic factors like BDNF (Bai *et al*, 2023; Luo *et al*, 2022). An intriguing aspect of our results is that this reduction in M1-associated markers was not accompanied by a corresponding increase in CD206, a commonly used marker of M2-like macrophages. One possible explanation lies in the role of extracellular matrix components such as CSPGs, which have been shown to constrain Microglia/Macrophage phenotypic conversion. In rodent SCI models, CSPGs abundant in the post-SCI scar, exert only modest effects on M1 cells but strongly prevent M2-like macrophages from maintaining a reparative state (Dyck *et al*., 2018; Francos-Quijorna *et al*., 2022). This blockade occurs through TLR4 signaling and is already evident by 7 days post-injury, the stage when inflammation would normally begin to resolve (Francos-Quijorna *et al*., 2022). Consequently, persistent CSPG accumulation interferes with the resolution phase and locks macrophages into a pro-inflammatory profile. Our data suggest that rMS effectively alleviates chronic M1-like neuroinflammation, which occurred alongside a trend toward reduced CSPG deposition. However, the absence of increased CD206 expression at 21 day post injury likely reflects the fact that the resolution window had already been disrupted earlier, such that M2-like populations could not expand despite reduced CSPG levels. Alternatively, it is possible that the magnitude of CSPG attenuation achieved in this study was insufficient to independently trigger a significant phenotypic shift toward an anti-inflammatory M2-like state at this chronic stage. Additionally, CD206 alone may not capture the full spectrum of reparative phenotypes, and future work including complementary markers may clarify these dynamics.

### Promotion of respiratory function through improved diaphragm activity

High cervical spinal cord injury (SCI) often results in significant disruption of descending respiratory pathways, leading to impaired neuromuscular control of the diaphragm and reduced respiratory efficiency (National Spinal Cord Injury Statistical, 2005; Warren *et al*, 2014; Winslow & Rozovsky, 2003). As expected, a significant reduction in VT (∼40%) was observed across all animals within the first 7 days post-injury, reflecting diminished neuromuscular drive stemming from phrenic pathway interruption, the present results suggest partial spontaneous recovery (25%) after 21 days post injury, although this increase was not sufficient to restore normal ventilation. By contrast, rMS-treated mice exhibited a significantly greater restoration of VT at 21 dpi, which reached almost complete recovery, indicating that rMS accelerated and amplified the limited spontaneous recovery.

During hypercapnia, which strongly activates chemoreceptor-driven respiratory drive and tests the ability to recruit ventilatory reserve, rMS animals exhibited stronger within-group recovery at D7 vs. D21, whereas sham-injured mice showed deficits in VT (and therefore VE). This suggests that rMS facilitates the recruitment of spared bulbospinal and interneuronal pathways under conditions of increased demand, thereby enhancing respiratory reserve capacity. These findings are consistent with previous reports showing that mid-cervical SCI impairs the ventilation during hypercapnic challenge, reflecting a loss of reserve capacity (Choi, Liao et al. 2005, Golder, Fuller et al. 2011). During normal poikilocapnic breathing, rMS produced a within-group recovery of VT and VE by day 21, whereas sham animals failed to restore VT and instead compensated by increasing breathing frequency. These divergent patterns suggest two distinct mechanisms that rMS enhanced inspiratory strength through greater phrenic motoneuron excitability, more effective motor-unit recruitment, and improved neuromuscular transmission, while sham animals relied on a less efficient “rapid shallow breathing” strategy to maintain ventilation. The absence of significant between-group differences likely reflects high variability in spontaneous breathing, yet the within-group recovery (D0 vs. D21) in rMS indicates a potential treatment effect.

Respiratory frequency (Bf) remained stable across time points and groups, indicating preserved central rhythm genic activity and suggesting that changes in VE were primarily driven by modulation of VT under 1% isoflurane anesthesia. Given the anesthetic suppression of accessory respiratory muscles, these findings suggest that the functional recovery mediated by rMS specifically arises from enhanced neural drive and mechanical output of the diaphragm. This finding is consistent with the idea that rMS may have triggered a long-term facilitation (LTF)-like effect that preferentially enhanced the amplitude of respiratory motor outputs rather than the central rhythm. Given its train-based, intermittent structure, our rMS protocol (9 trains with inter-train pauses) plausibly fulfils these temporal requirements, and previous work has likewise shown that rMS can activate BDNF/ERK signaling cascades (Wang, Crupi et al. 2011, Peng, Zhou et al. 2018). In support of this interpretation, spinal iTBS has been reported to increase diaphragm EMG burst amplitude for periods extending beyond the stimulation session, indicating a sustained enhancement of inspiratory drive at the motoneuron level (Lee and Vinit 2024). Thus, the absence of frequency modulation in our rMS data likely reflects a mechanism in which spinal plasticity strengthens inspiratory force generation without altering the central rhythm generator.

The Diaphragm activity recovery found in our study is in line with previous evidences that magnetic stimulation can restore inspiratory drive, increase diaphragm excitability, and induce neuroplastic changes in descending respiratory circuits (Lee & Vinit, 2024; Michel-Flutot *et al*, 2022; Randelman *et al*, 2021). At the same time, the observed trend toward reduced CSPG levelswas associated with a trend toward an increased number of labeled neurons at the level of the lesion, where phrenic motoneuron are located. Thus, one hypothesis is that the combination of rMS-induced trends in CSPG attenuation and enhanced neuronal survival may have strengthened spared motoneuron and connections at the level of the PMN pool and around.

Beyond reducing inhibitory CSPGs and enhancing neuronal survival, additional evidence indicates that rMS/rTMS promotes regenerative plasticity in spared phrenic motoneurons and their descending inputs. In C2 hemisection rats, high-frequency rTMS increased GAP-43 expression in ventrolateral cervical tracts, consistent with axonal sprouting within corticospinal-phrenic pathways (Michel-Flutot *et al*., 2022). Similar upregulation of GAP-43 has been reported after iTBS in incomplete compression SCI models (Marufa *et al*, 2021) and trans-spinal iTBS has been shown to augment inspiratory drive may via enhanced synaptic activity within the phrenic nucleus and likely modulation of excitatory and inhibitory inputs to phrenic motoneurons in a C3 contusion model (Lee & Vinit, 2024). Together, these mechanisms together are likely to contribute to the improved functional outcomes seen in rMS-treated animals. Similarly, previous work showed that rTSMS minimizes cavity formation and preserves axons in both contusion and transection models, resulting in better locomotor recovery (Bai *et al*., 2023). Such mechanistic insights will be essential to refine stimulation protocols and enhance their translational relevance.

### Clinical Relevance

These findings have clear translational relevance. Non-invasive rMS is already FDA-approved for the treatment of depression and has demonstrated safety and efficacy in managing spasticity and neuropathic pain in SCI patients (Benavides *et al*, 2025; Cotovio *et al*., 2023). Our data suggest that rMS could be repurposed to directly target the spinal cord injury site. By reducing glial scarring, mitigating neuroinflammation, and enhancing neuroprotection and plasticity, rMS may synergize with rehabilitative therapies to accelerate functional recovery. Importantly, as a non-invasive and repeatable modality, rMS presents minimal clinical risk and strong therapeutic potential. These results warrant further clinical exploration and may pave the way toward incorporating rMS as a standard adjunct in SCI recovery protocols.

### Limitations and Strengths

One limitation of the present study is the use of a single model of cervical spinal cord injury, C3 hemi-contusion, which may not fully reflect the heterogeneity and complexity of human cervical SCI. Different injury types, severities, and segmental levels may engage distinct respiratory and inflammatory responses, potentially affecting the generalizability of our findings. However, the C3HC model is relatively recent and less widely used but clinically relevant paradigm that closely mimics many key features of incomplete cervical SCI in humans, including partial preservation of descending motor pathways and compromise in ventilatory capacity.. Its reproducibility and translational fidelity make it a suitable starting point for evaluating the therapeutic potential of rMS in modulating respiratory recovery. Another limitation lies in the temporal resolution of the respiratory assessments, which were conducted at three discrete time points, baseline, 7 days post-injury, and 21 days post-injury. While these time points were strategically selected to represent the pre-injury state, the acute phase of injury, and the early subacute recovery phase, they may miss transient or delayed neuroplastic changes occurring outside this window. Nonetheless, this focused time-course design enabled clear comparisons across critical stages of injury and recovery while minimizing the impact of biological variability. Importantly, the observed effects of rMS on both ventilatory parameters and diaphragm activity were robust despite the limited temporal sampling (Michel-Flutot *et al*., 2022). A critical area for future exploration lies in the optimization of rMS parameters, including frequency, burst pattern, and stimulation intensity. While our standardized 10 Hz protocol elicited functional gains, the field lacks a comprehensive “dose-response” map for respiratory recovery. Different frequencies may engage distinct molecular pathways. For instance, intermittent theta burst stimulation (iTBS) is often associated with more potent long-term potentiation (LTP)-like effects via rapid BDNF-TrkB signaling, whereas continuous low-frequency stimulation might more effectively modulate homeostatic plasticity or suppress hyperexcitability. Finally, regarding treatment duration, our short-term rMS regimen elicited significant improvements in tidal volume, respiratory reserve, and diaphragm electromyographic output by Day 21 suggests a strong capacity for promoting early-phase neuroplasticity. This aligns with recent evidence f that extended stimulation protocols do not necessarily yield superior outcomes (Lee & Vinit, 2024). Similarly, comparative studies demonstrating that a two-week intervention was sufficient to reach a therapeutic plateau in respiratory recovery, with a four-week regimen offering no additional functional gain (Michel-Flutot *et al*., 2022). These findings support the notion that even limited, subacute stimulation protocols can yield meaningful functional gains and may serve as an efficient entry point for optimizing stimulation parameters in future translational studies.

### Conclusion

This study provides converging evidence that targeted repetitive magnetic stimulation (rMS) applied to the injured cervical spinal cord markedly enhances respiratory recovery after high cervical SCI. rMS-treated animals recovered larger tidal volumes and minute ventilation across awake and challenge conditions, and exhibited stronger diaphragm activity on EMG in both hemidiaphragms. These functional gains were paralleled by dramatic changes in the injury microenvironment. rMS reduced indicators of reactive astrogliosis (GFAP) and fibrotic scarring (PDGFRβ), and lowered pro-inflammatory microglial activation (Iba1, CD68), without affecting CD206+ cell levels. Neuronal survival was improved (more NeuN+ cells), and inhibitory extracellular matrix (PNNs) tends to diminish. Thus, rMS appears to relieve multiple barriers to plasticity, creating a pro-regenerative milieu. These findings are consistent with prior reports that high-frequency magnetic stimulation can suppress astrocyte and microglial reactivity and inflammation, reduce glial scar formation, and support axonal preservation. Importantly, our results highlight the translational promise of spinal rMS as a non-invasive neuromodulatory therapy for cervical SCI. By attenuating scar and inflammation while strengthening the spared MN and connections at the level of the PMN pool and around, rMS effectively accelerated and amplified the limited spontaneous recovery. In summary, targeted rMS significantly promotes ventilatory function and neural plasticity after cervical SCI, underscoring its potential as a therapeutic strategy. Future studies should optimize stimulation protocols a nd explore combination with rehabilitation to maximize respiratory recovery, with the ultimate goal of improving breathing outcomes for individuals with cervical SCI.

## MATERIALS AND METHODS

### Animal Care and Use Statement

The animal protocol was designed to minimize pain or discomfort to the animals. All experimental procedures were in accordance with the European Community guiding principles on the care and use of animals (EU Directive 2010/63/EU), and approved by the Ethics Committee Charles Darwin CEEACD/N 5 (Project authorization APAFIS No. 201901301500576).

### Experimental Design

A total of 29 adult male Swiss mice (5 weeks old) were used to investigate the effects of repetitive trans-spinal magnetic stimulation (rMS) on respiratory function after cervical spinal cord injury. Mice were randomly assigned to four experimental groups: (1) laminectomy + sham rMS (n = 4); (2) laminectomy + rMS (n = 4); (3) C3 hemi-contusion (C3HC) + sham rMS (n =10); and (4) C3HC + rMS (n =11). Respiratory function was assessed using whole-body plethysmography at baseline, post-injury, and after the rMS intervention, with tidal volume (Vt) and respiratory frequency (Bf) recorded. Diaphragm activity was further evaluated through in-situ electromyography (EMG) under terminal anesthesia. Spinal cord tissues were harvested for immunofluorescence to assess neuroinflammatory and structural markers. rMS or sham stimulation was applied once daily for two weeks starting 7 days post-surgery.

### Surgical procedures

Animals were placed in a closed chamber for anesthesia induction with isoflurane (4%) and maintained throughout the procedure with a facial mask (1.5-2.5% isoflurane) in 100% O2. The dorsal skin and underlying muscles above the second cervical up to the first thoracic vertebrae was retracted. A dorsal laminectomy were performed at C3/C4 to expose the spinal cord. A precision Impactor Device (RWD life science; 68,099 II) with a 1.5 mm tip impactor will be used to perform the C3/4 hemi-contusion. The impactor parameters are set-up as follows: a mean depth of 2.6 ± 0.41 mm, a speed of 1.0 ± 0.1 m/s and a dwell time of 0.60 ±0.01 s (mean ± SEM). After contusion, the wounds and skin were closed. The isoflurane vaporizer was turned off, and the mice received subcutaneous injections of sulfadoxine (Borgal, 0.2 mL per mouse; 0.1 mL diluted in 1 mL NaCl) and buprenorphine (Buprécare, 0.5 mL per mouse; 0.05 mL diluted in 1 mL NaCl). After surgery, the animals were placed on a heated pad to recover. The animals were then placed in a cage containing water and a recovery diet gel, both in a small Petri dish, and then solid food were provided in the cage. Food and water were also provided on the top grid of the cage. Animals that showed a reduction of more than 20% tidal volume (Vt) from baseline were divided into two groups: (1) C3HC + sham rMS (n = 7); (2) C3HC + rMS (n = 10).

### Repetitive Magnetic Stimulation (rMS) Protocol

rMS protocol was performed using the magnetic stimulator MAGPRO R30 (Magventure, Farum, Denmark) connected to a figure-of-eight coil (Cool-B65), delivering a unique biphasic pulse with the intensity of the stimulus expressed as a percentage of a maximum output of the stimulator (% MO). The protocol (10 Hz, 9 trains of 100 biphasic pulses, separated by 30 s intervals between trains delivered at 80% MO, 900 stimulations per protocol) was applied in awake restrained animals. This protocol induced a long-lasting increase in phrenic excitability in anesthetized, intact rats(Michel-Flutot *et al*., 2022). Control animals received a Sham rMS protocol (e.g., no stimulation but the same time spent in the custom-designed restraining device, Figure 2). This rMS protocol was applied 7 days postinjury for 2 weeks (once a day, 5 days per week) (Figure 2).

### Breathing Recording Using Plethysmography

Respiratory function was assessed via whole-body plethysmography (Emka, France) at baseline, 7 days post-surgery, and following rMS treatment (Day 21). The mice were placed in a hermetically sealed chamber (comprising a 250 mL animal compartment and a 150 mL reference chamber) under a continuous 0.5 L/min bias flow. Subsequently mice were exposed to the following test gases in chronological order: 5 min 1% Isoflurane, 5 min 5% CO_2_, 5 min 1% Isoflurane, 10 min ambient air. This phase limit stress and resulted in calm animals, thereby allowing successful recordings. Plethysmography signals were digitized and analyzed using LabChart 8 Pro software (ADInstruments, Dunedin, New Zealand). Quantified ventilatory parameters included tidal volume (VT), breathing frequency (Bf), minute ventilation (VE), and inspiratory/expiratory times (Ti and Te).

### Electrophysiological Recording of the Diaphragm

After 21 days post-rMS treatment, terminal diaphragmatic electromyography (dia-EMG) were performed to evaluate respiratory motor output. Mice were placed in a supine position on a regulated heating pad to maintain a constant core body temperature (37.5 ± 1 °C). Following a midline laparotomy to expose the diaphragm, dia-EMG activity was recorded from the ventral, medial, and dorsal regions of both the ipsilateral and contralateral hemi-diaphragms during spontaneous breathing. Signals were obtained using custom-made bipolar silver surface electrodes positioned on the muscle fibers. The dia-EMG signals were amplified (Model 1800; gain: 100; A-M Systems, Everett, WA, USA) and bandpass-filtered (100 Hz – 10 kHz). Data were digitized using a PowerLab acquisition system (Acquisition rate: 4 k/s; ADInstruments, Dunedin, New Zealand). For quantitative analysis, raw signals were rectified and double-integrated (50 ms decay constant) using LabChart 8 Pro software (ADInstruments). The amplitude of at least 10 double-integrated diaphragmatic EMG inspiratory bursts during normoxia was calculated for each animal from the injured and the intact sides. After the experiment, the animals will be euthanized by exsanguination, followed by intracardiac perfusion of 4% paraformaldehyde (4°C) for tissue fixation and subsequent harvesting.

### Tissue Processing

After fixation, the C1-C8 segment of the spinal cord will be dissected and immediately placed in cold 4% paraformaldehyde (P6148 Sigma-Aldrich, Darmstadt, Germany) for 24 h and then cryoprotect in 30% sucrose (in 0.9% NaCl, S9888, Sigma-Aldrich, Darmstadt, Germany) for 48 h and store at −80°C. Frozen transversal (C1- C8 spinal cord) free-floating sections (30 μm) will be obtained using a Thermo Fisher CryoStar NX70 cryostat. Spinal cord sections will be stored in a cryoprotectant solution (sucrose 30% (pharma grade, 141621, AppliChem, Darmstadt, Germany), ethylene glycol 30% (BP230-4, Fisher Scientific, Illkirch, France) and polyvinylpyrrolidone 40 (PVP40-100G; Sigma-Aldrich 1%) in phosphate-buffered saline (PBS) 1X (BP665-1; Fisher Scientific, Illkirch, France) at −22°C. Every fifth section from C1- C8 will be used for lesion evaluation to examine the extent of injury using cresyl violet histochemistry: 30 min in cresyl violet solution (0.001% cresyl violet acetate (C5042-10G, Sigma-Aldrich, Darmstadt, Germany) and 0.125% glacial acetic acid (A/0400/PB15, Fisher Scientific, Illkirch, France) in distilled water), 30 s in 70% ethanol, 30 s in 95% ethanol, 2 × 10 s in 100% ethanol (E/0600DF/17, Fisher Scientific, Illkirch, France) and 2 min in xylene (X/0100/PB17, Fischer Scientific, Illkirch, France). Then, they will be coverslipped using Eukitt® mounting medium, and slide microphotographs will be taken with a slide scanner (Aperio AT2, Leica, France). Each injury was then digitized and analyzed with ImageJ 1.54k software (National Institutes of Health, Bethesda, MD, USA; Schneider et al., 2012). The extent of the injury on the injured side was calculated using a reference to a complete hemicontusion (which is 100% of the hemicord).

### Immunohistochemistry

Several markers of interest were used on spinal cord samples from C3HC for qualitative assessments. Free-floating transverse sections of the C1-C4 and C5-C8 spinal cord stored in cryoprotectant solution have been washed 3 times in PBS 1X and placed in blocking solution (normal donkey serum (NDS) 5% and 0.2% Triton 100X in PBS 1X) for 30 min at room temperature. For primary labeling, sections were incubated overnight on an orbital shaker at 4 °C. The following primary antibodies and/or biotinylated lectins were used: PDGFRβ (Abcam ab32570, 1/1000, rabbit polyclonal), GFAP (PA5-18598, 1/1000, goat polyclonal), Iba1 (Abcam ab5076, 1/400, goat polyclonal), CD68 (ThermoFisher, 14-0681-82, 1/100, rat polyclonal), CD206 (ThermoFisher, MA5-16869, 1/100, rat polyclonal), Wisteria Floribunda Lectin (WFA, Vector laboratories, Les Ulis, 1/2000) was used to labeled chondroitin sulfate proteoglycans (CSPGs). After 3 times PBS 1X washes, sections were incubated for 2 h at room temperature with the appropriate secondary reagents. Antibodies were visualized using Alexa Fluor 647 Donkey anti-rabbit (ThermoFisher, A-31573, 1/1000), Alexa Fluor® 594 donkey anti-goat (ThermoFisher, A-11058, 1/1000), Alexa Fluor 647 Donkey anti-rat (ThermoFisher, A78947, 1/1000), biotinylated WFA binding was detected using Alexa Fluor 488 Avidin (Molecular Probes, Illkirch, 1/1000) and they were washed again 3 times with PBS 1X. Sections were incubated with NeuroTrace™435/455 (ThermoFisher, N21479, 1/1000), a neuronal marker (Nissl stain), for 10 min and then wash again 3 times with PBS 1X. Images of the different sections will be captured with a Hamamatsu ORCA-R2 camera mounted on an Olympus IX83 P2ZF scanning microscope.

### Data Processing and Statistical Analyses

All data are presented as mean ± standard deviation (SD). Statistical analyses were performed using GraphPad Prism (version 10.1.2, GraphPad Software, USA). The normality of data distributions was assessed using the Shapiro-Wilk test. A significance threshold of p < 0.05 was applied throughout. Comparisons between two groups (e.g., sham rMS vs. rMS) for continuous variables such as body weight, lesion extent, diaphragmatic EMG activity, respiratory parameters, and immune marker quantifications were analyzed using unpaired t-test for normally distributed data, or the Mann-Whitney test for non-parametric data. For comparisons among more than two independent groups (e.g., laminectomy + sham rMS, laminectomy + rMS, C3HC + sham rMS, C3HC + rMS), one-way ANOVA followed by Tukey’s multiple comparisons test was used for parametric data. For non-parametric data, the Kruskal-Wallis test followed by Dunn’s post hoc test was applied. Repeated measures two-way ANOVA was used to assess the effects of treatment (sham rMS vs. rMS) over time (D0, D7, D21) for longitudinal variables such as respiratory measurements. Multiple comparisons were corrected using the Šidák or Tukey method. Sample sizes and statistical test details are reported in each figure legend.

## Author contributions

Conceptualization: WC, SV, IV

Methodology: WC, SV

Investigation: WC, SV, IV

Visualization: WC

Funding acquisition: WC, SV

Project administration: WC, SV, IV

Resources: SV, IV

Supervision: SV, IV

Writing – original draft: WC

Writing – review & editing: WC, SV, IV

## Data and materials availability

All data needed to evaluate the conclusions in the paper are present in the paper and/or the Supplementary Materials.

