## Supplementary Materials for "Cervical Repetitive Magnetic Stimulation Enhances Respiratory Recovery by Modulating Neuronal Plasticity After Cervical Spinal Cord Injury"

SUPPLEMENTARY MATERIAL

Supplementary Figures

Legends for Supplementary Data

Tables

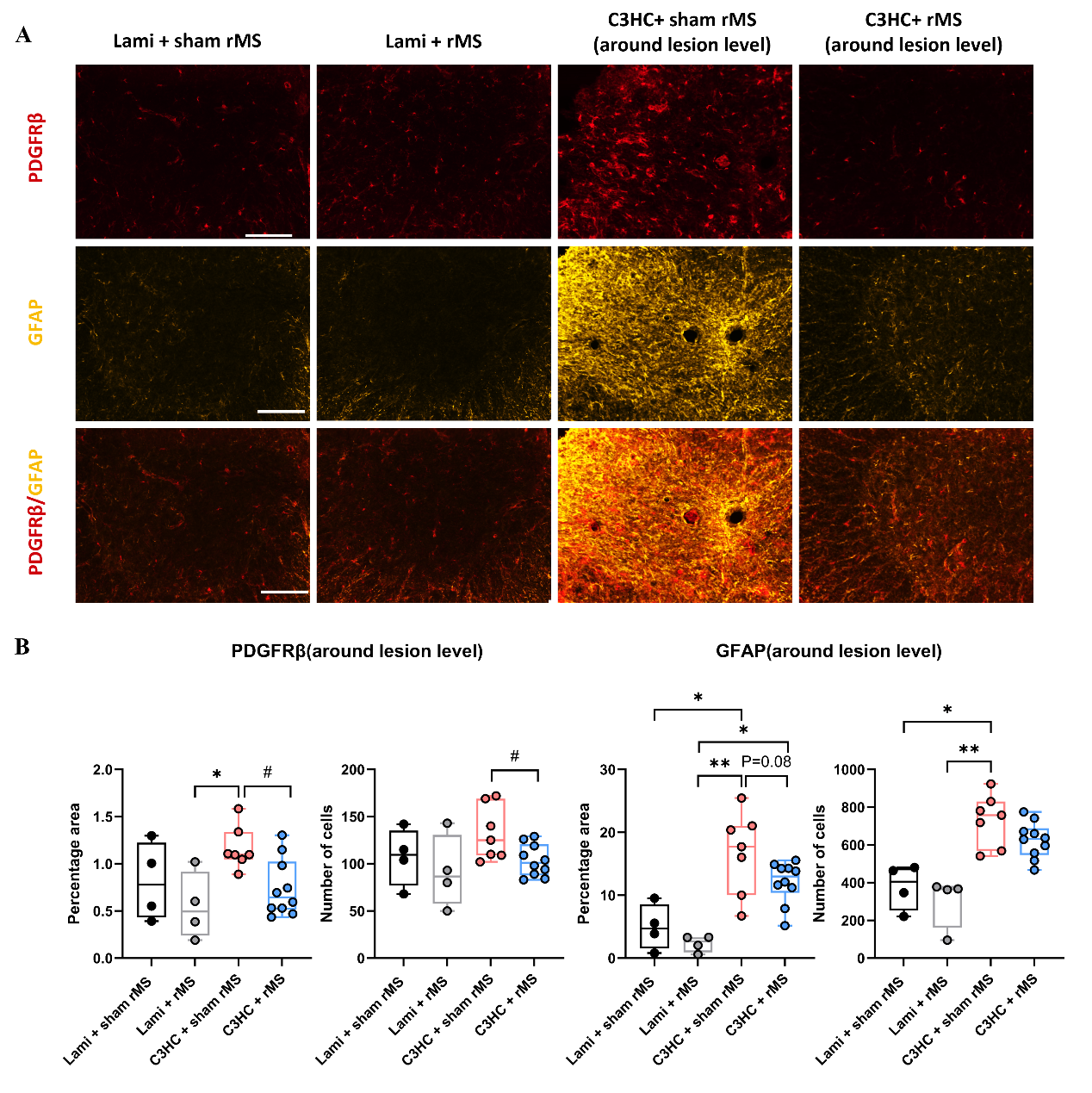

**Supplementary Figure 1. PDGFRβ and GFAP immunoreactivity at 21 days post-injury in the cervical spinal cord around lesion level.**

**(A)** Representative immunofluorescence images showing PDGFRβ (red), GFAP (yellow), and merged channels around the lesion area of Laminectomy + sham rMS, Laminectomy + rMS, C3HC + Sham rMS, and C3HC + rMS-treated animals at 21 days post-injury. Scale bar: 500 µm. **(B)** Quantification of PDGFRβ+ percentage area, GFAP+ percentage area, and cell number around the lesion site at the ventral part of spinal cord. Each dot represents one animal. Histograms (Min to max, Box and Whiskers) show data distribution across the four groups. Statistical significance across the four experimental groups was determined using a Kruskal-Wallis test followed by Dunn’s post-hoc multiple comparisons test (*) *p<0.05, **p<0.01, while direct comparisons between the C3HC + sham rMS and C3HC + rMS-treated groups were performed using a Mann-Whitney tes (^#^) ^#^p<0.05.

In the peri-lesional region, PDGFRβ and GFAP were co-expressed within the fibrotic and astroglial scar. Mice receiving rMS after C3HC exhibited a reduction in PDGFRβ immunoreactivity compared to the sham-stimulated group, both in percentage area (1.17 ± 0.23 % vs. 0.74 ± 0.30 %, *p* < 0.05) and in cell number (132 ± 29 vs. 104 ± 17, *p* < 0.05). Similarly, GFAP-positive area decreased significantly following rMS compared to sham rMS (16.77 ± 6.51 % vs. 12.09 ± 3.32%, p = 0.08), along with a lower number of GFAP-positive cells but without significant difference (732 ± 137 vs. 625 ± 95).

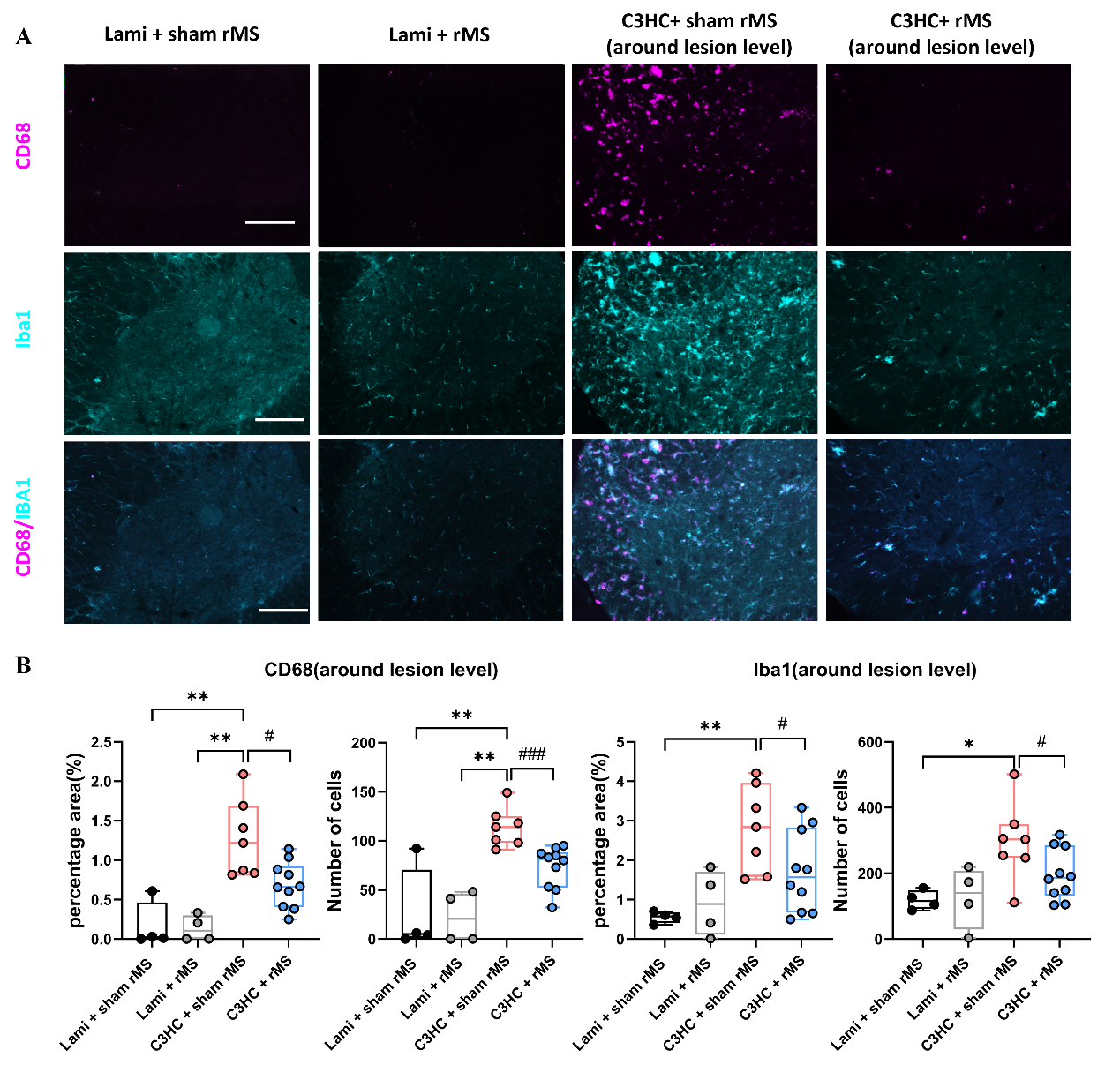

**Supplementary Figure 2. CD68 and Iba1 immunoreactivity at 21 days post-injury in the cervical spinal cord around lesion level.**

**(A)** Representative immunofluorescence images showing CD68 (purple), Iba1 (cyan), and merged channels around the lesion area of Laminectomy + sham rMS, Laminectomy + rMS, C3HC + Sham rMS, and C3HC + rMS-treated animals at 21 days post-injury. Scale bar: 500 µm. **(B)** Quantification of CD68+ percentage area, Iba1+ percentage area, and cell number around the lesion site at the ventral part of spinal cord. Each dot represents one animal. Histograms (Min to max, Box and Whiskers) show data distribution across the four groups. Statistical significance across the four experimental groups was determined using a Kruskal-Wallis test followed by Dunn’s post-hoc multiple comparisons test (*) *p<0.05, **p<0.01, ***p<0.001, while direct comparisons between the C3HC + sham rMS and C3HC + rMS-treated groups were performed using a Mann-Whitney test (^#^) ^#^p<0.05, ^##^p<0.01, ^###^p<0.001.

Iba1 and CD68, co-expressed by pro-inflammatory microglia/macrophages, also showed significant reduction after rMS. Iba1-positive area decreased compared to sham rMS group (2.80 ± 1.09 % to 1.70 ± 1.02 %, p < 0.05), and the number of Iba1-positive cells also decreased (296 ± 118 to 196 ± 77, p < 0.05). CD68-positive area was also significantly reduced (1.28 ± 0.49 % vs. 0.68 ± 0.29 %, p < 0.01), as was the number of CD68-positive cells (114 ± 20 vs. 73 ± 21, p < 0.01).

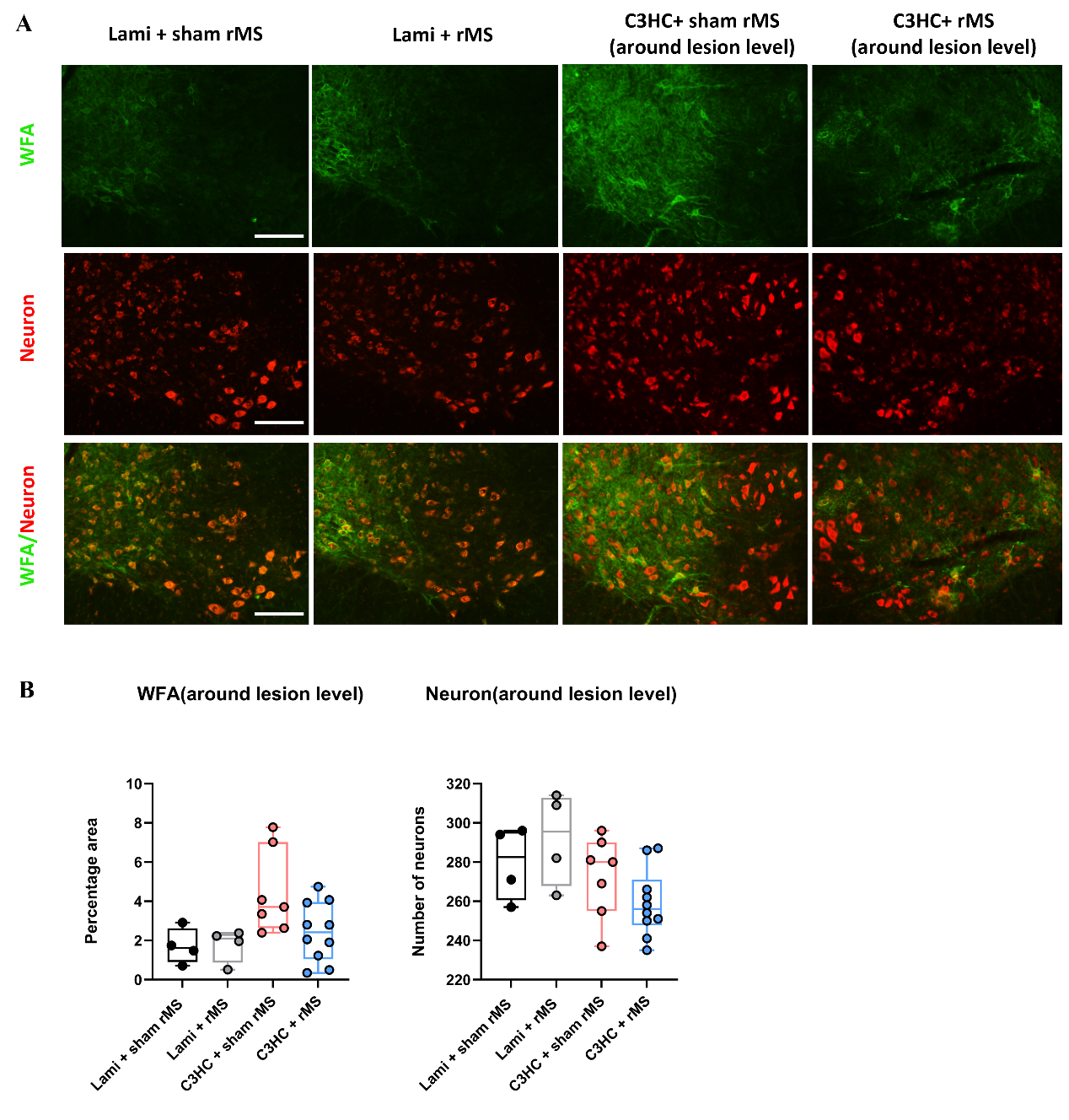

**Supplementary Figure 3. WFA staining and ventral neuron changes at 21 days post-injury in the cervical spinal cord around lesion level.**

**(A)** Representative immunofluorescence images showing WFA (green), neuronal marker (red), and merged channels around the lesion area of Laminectomy + sham rMS, Laminectomy + rMS, C3HC + sham rMS, and C3HC + rMS-treated animals at 21 days post-injury. Scale bar: 500 µm. **(B)** Quantification of WFA-positive percentage area, number of ventral neurons around the lesion site. Each dot represents one animal. Histograms (Min to max, Box and Whiskers) show data distribution across the four groups. Statistical comparisons across the four experimental groups were performed using a Kruskal-Wallis test followed by Dunn’s post-hoc multiple comparisons test, while direct comparisons between the C3HC + sham rMS and C3HC + rMS-treated groups were conducted via a Mann-Whitney test; no statistically significant differences were detected among the groups.

WFA staining, used to assess perineuronal net density, was lower in the rMS group compared to sham but without difference (4.42 ± 2.12 % vs. 2.44 ± 1.51), suggesting a reduction in extracellular matrix components. NeuN were assessed in the ventral part, with no significant difference between sham rMS and rMS groups (273 ± 21 vs. 259 ± 17).

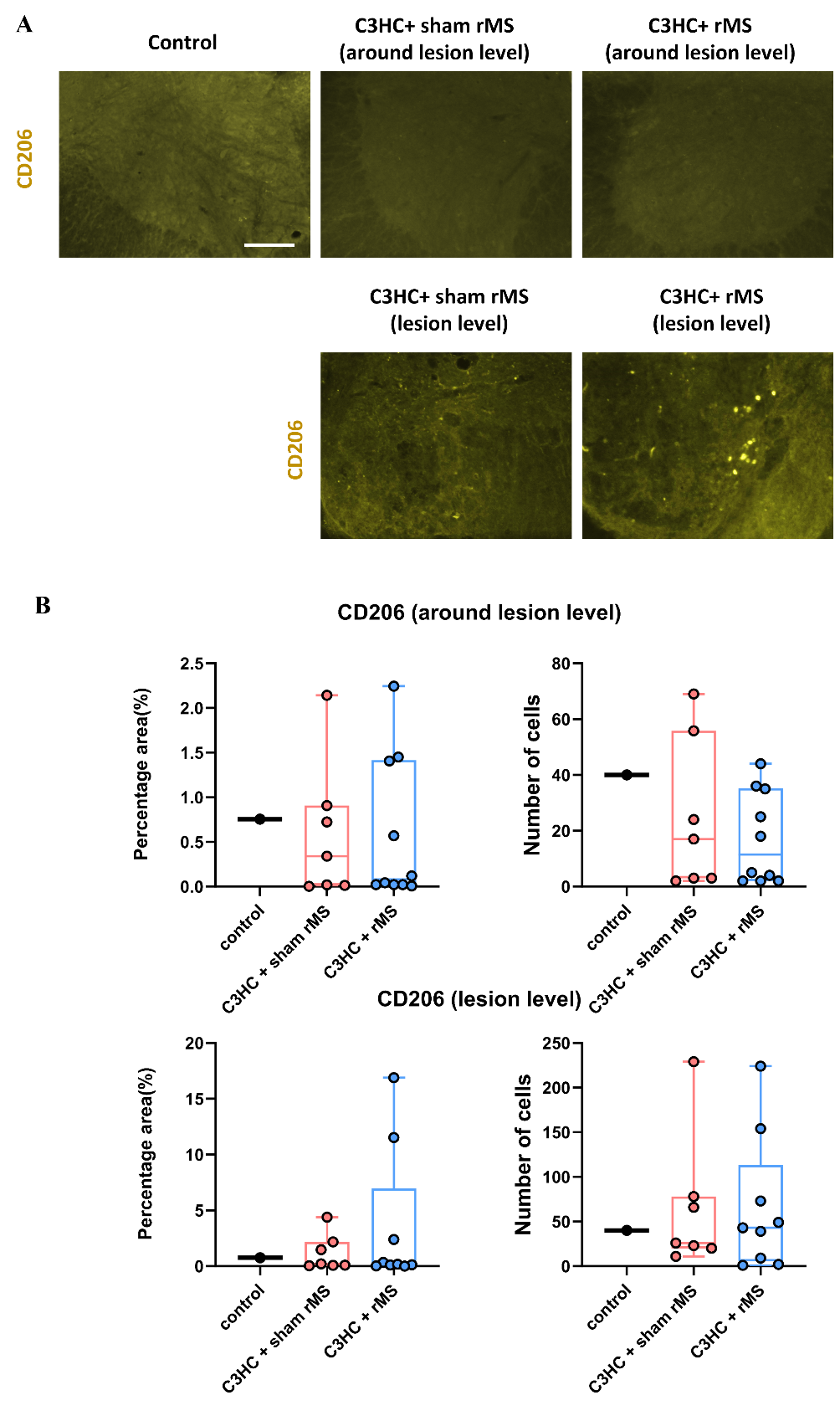

**Supplementary Figure 4. CD206 staining at 21 days post-injury in the cervical spinal cord at and around lesion level.**

**(A)** Representative immunofluorescence images showing CD206(yellow) at and around the lesion area of control, C3HC + sham rMS, and C3HC + rMS-treated animals at 21 days post-injury. Note: The “Control” group represents naive mice that underwent no surgical or anesthetic procedures. Scale bar: 500 µm. **(B)** Quantification of CD206-positive percentage area at and around the lesion site. Each dot represents one animal. Histograms (Min to max, Box and Whiskers) show data distribution across the four groups. Statistical comparisons among the three groups were performed using a Kruskal-Wallis test; no significant differences were observed between any of the groups.

CD206 showed no significant differences between groups in percentage area (0.76 ± 0.0 vs. 0.59 ± 0.77 % vs. 0.59 ± 0.81 %, n.s.) or cell number (20 ± 1 vs. 25 ± 27 vs. 17 ± 17, n.s.) around lesion level or in in percentage area (0.76 ± 0.0 vs. 1.2 ± 1.6 % vs. 3.5 ± 1.3 %, n.s.) or cell number (20 ± 1 vs. 65 ± 77 vs. 66 ± 76, n.s.) at lesion level.

Table 1. Respiratory parameters and body weight at different timepoints after SCI and treatment

| **Group** | **Timepoint** | **Weight**  **(g)** | **VT**  **(µL/g)** | **Bf (breaths/min)** | **VE (mL/min)** |
| --- | --- | --- | --- | --- | --- |
| Laminectomy + Sham rMS  (n = 4) | D0 | 39 ± 3 | 5.08 ± 1.08 | 164 ± 20 | 32.21 ± 8.25 |
|  | D7 | 39 ± 4 | 6.35 ± 0.50 | 167 ± 22 | 41.69 ± 7.12 |
|  | D21 | 39 ± 4 | 7.34 ± 1.74 | 164 ± 16 | 47.41 ± 13.38 |
| Laminectomy + rMS  (n = 4) | D0 | 38 ± 4 | 5.86 ± 1.00 | 169 ± 25 | 37.57 ± 7.20 |
|  | D7 | 40 ± 4 | 6.61 ± 1.89 | 164 ± 28 | 43.75 ± 18.97 |
|  | D21 | 40 ± 6 | 7.03 ± 2.49 | 157 ± 27 | 43.41 ± 13.38 |
| C3HC + Sham rMS  (n = 8) | D0 | 38 ± 2 | 5.81 ± 0.88 | 176 ± 27 | 38.62 ± 6.87 |
|  | D7 | 36 ± 3 | 3.08 ± 0.88 | 131 ± 53 | 15.63 ± 7.84 |
|  | D21 | 37 ± 2 | 3.87 ± 1.01 | 152 ± 16 | 21.94 ± 6.48 |
| C3HC + rMS  (n = 10) | D0 | 39 ± 2 | 5.71 ± 0.61 | 171 ± 33 | 37.95 ± 9.48 |
|  | D7 | 34 ± 3 | 3.88 ± 1.01 | 145 ± 56 | 20.52 ± 10.45 |
|  | D21 | 36 ± 3 | 6.02 ± 1.15 | 167 ± 39 | 36.53 ± 11.56 |
